# Effect of sleep stages on patterns of fNIRS hemodynamic response to auditory paradigms in one-month-old Gambian and UK infants

**DOI:** 10.1101/2025.07.28.667238

**Authors:** Maria Rozhko, Merel van der Straaten, Borja Blanco, Isobel Greenhalgh, Johann Benerradi, Sophie E. Moore, Clare E. Elwell, Sarah Lloyd-Fox, Anna Blasi, the BRIGHT Project Team

## Abstract

**Significance:** Functional near-infrared spectroscopy (fNIRS) has advanced our understanding of early brain development, especially infant responses to social and auditory stimuli. Unlike older children and adults, very young infants are often assessed during natural sleep to reduce movement and ensure sufficient data quality. Yet the impact of sleep stage on fNIRS signals and how it might affect interpretations of early brain activation patterns remains unclear.

**Aim:** This study investigates the effect of sleep stages on fNIRS-measured hemodynamic responses to two auditory paradigms across different global populations of one-month-old infants.

**Approach:** In total, 46 Gambian and 40 UK infants in quiet or active sleep were tested using (1) social selectivity and (2) a habituation and novelty detection paradigms.

**Results:** In the UK cohort, active sleep was associated with a stronger initial response and greater habituation compared to quiet sleep. In contrast, Gambian infants in quiet sleep showed more widespread activation and evidence of habituation, while infants in active sleep showed no habituation. No sleep stage effects were observed for response in the social selectivity paradigm in either group.

**Conclusions:** Different effects of sleep stages were observed across the two cohorts and paradigms and should be carefully considered in neuroimaging studies.

## 1 Introduction

The early postnatal period represents a critical time for studying brain development, marked by intense neural growth and reorganization, including synapse formation and pruning, glial proliferation, and myelination, shaped by both genetic programming and environmental experiences.^1^ Several neuroimaging methods, including electroencephalography (EEG), magnetoencephalography (MEG), functional magnetic resonance imaging (fMRI), and functional near-infrared spectroscopy (fNIRS), are effective for investigating brain function in neonates and infants. While EEG and MEG provide direct measures of neural activity, fMRI and fNIRS offer indirect measures via hemodynamic responses associated with underlying neural activity. Among these, fNIRS stands out as the most portable, cost-effective, and least susceptible to movement artifacts, making it particularly well-suited for research outside specialized neuroimaging laboratories.^2^ By measuring hemodynamic responses through changes in oxyhemoglobin (HbO) and deoxyhemoglobin (HbR) signals, fNIRS enables functional characterization of cortical regions.^3^

Despite its higher tolerance to movement, acquiring brain imaging data on awake infants in the first weeks of life remains challenging due to limited quiet alertness, frequent movements, and brief visual attention at this age. As a result, fNIRS paradigms often focus on measuring brain responses to auditory stimuli and task-free (or resting-state) brain activity during natural sleep. However, sleep is not a static state: in infant studies, recordings collected during sleep may span one or more cycles of active sleep (AS), a precursor to rapid eye movement (REM) sleep, which is associated with synapse formation and is characterized by increased activity in the sensorimotor region; or quiet sleep (QS), a precursor to non-rapid eye movement (NREM) sleep, which is linked to synaptic pruning and network refinement.^4^

Sleep stages also differ in how they manifest behaviorally. AS is marked by irregular breathing, rapid eye movements, facial muscle twitches, sucking, and startles, while QS involves regular breathing, minimal movement of the face and body, and no eye movements.^5^ In addition, infants are more reactive to external stimuli in AS, while QS is associated with delayed responses and a reduced likelihood of arousal.^6^ These protective qualities of QS are believed to be present from 33-34 weeks of gestation.^7^

Building on the behavioral differences between QS and AS described above, several studies have examined whether sleep stages are also associated with distinct brain responses to sensory stimuli in infants, predominantly using techniques such as EEG and MEG. First, studies have investigated whether the amplitude of neural responses or the proportion of infants showing a significant response differ by sleep stage. These studies have consistently found stronger responses in QS than in AS.^8–11^ For example, a study with 20 full-term neonates at two days of age measured somatosensory evoked potentials using EEG in response to wrist nerve stimulation and found that AS and wakefulness were associated with lower-amplitude neural responses compared to QS.^8^ The same was shown to be true for magnetic fields evoked by auditory speech stimuli, which were also lower in amplitude in AS than in QS, in a group of ten^10^ and six^11, 12^ newborns. Another study examined somatosensory responses to tactile finger stimulation with MEG in 46 full-term neonates 1 - 23 days of age and found that a significant response was present in 90% of QS infants compared to only 50% of AS infants, indicating a more robust response in QS than in AS.^9^

Second, studies have examined the effect of sleep stage on stimulus discrimination (i.e., the brain’s ability to detect a change in stimulus features). EEG studies on auditory change detection in infants have produced mixed findings, with some suggesting that (a) in neonates, the change detection response is present in AS and QS and the amplitude of response to standard and deviant stimuli does not significantly vary according to sleep stage,^12–14^ (b) the change detection response is stronger (i.e. there is larger difference between response amplitude to standard and deviant sounds) in AS compared to QS in neonates^15^ and 2-to-3-month olds,^16^ or, on the contrary, (c) the response is stronger in QS compared to AS in 2-month old infants.^17^

Third, a smaller number of studies have explored how sleep stage influences habituation responses, specifically, the presence, strength, and timing of neural habituation to repeated stimuli.^18, 19^ Based on the evidence of an attenuated and delayed behavioral response to external stimuli in QS compared to AS,^6^ it would be reasonable to expect a faster neural habituation (gradual reduction in response to repeated stimuli) in QS. However, in an EEG study with 9.4±1.2-weekold infants McNamara et al. showed that while habituation to tactile stimulation was evident in both AS and QS, AS was associated with a more rapid habituation compared to QS.^18^ Similarly, in another study that measured sensory gating - a decrease in EEG response amplitude between two subsequent presentations of an auditory stimulus - at three months and four years of age, a significant reduction of response was present in AS, but not QS for both age groups. The latency of response was found to be longer in QS compared to AS for the infants, but not for four year olds.^19^

The hemodynamic response to auditory stimuli, discrimination between auditory stimuli, and habituation have previously been studied using fNIRS in sleeping newborns^20–28^ and infants aged 2-13 months.^29–33^ Among these, only two studies included sleep stage as a factor, neither of which studied habituation or auditory discrimination. Kotilahti et al. examined the effect of sleep stages on the location, amplitude, and latency of peak (time until the largest change in HbO and HbR concentration from baseline) of hemodynamic responses to simple sinusoidal beep stimuli in 20 full-term 1-3-day-old neonates. Contrary to the results of the EEG and MEG studies, they identified higher response amplitude in AS compared to QS, but there were no differences in the latency or the cortical location of the response between sleep stages.^25^ In a study that investigated response to speech and music sounds in 13 full-term 1-4-day-old neonates using fNIRS, no sleep stage effects were found.^26^

In summary, evidence from the EEG and MEG literature suggests that QS is associated with stronger neural responses to sensory stimuli, while AS may support faster or more dynamic processing, such as sensory gating and habituation. However, the limited evidence from the fNIRS studies suggests either greater hemodynamic response in AS compared to QS or no difference in response according to sleep stage. To our knowledge, no studies have examined the effect of sleep stage on auditory discrimination or habituation measured using fNIRS in young infants. Therefore, the evidence remains mixed and potentially paradigm or modality dependent, highlighting the need to systematically examine how sleep stage modulates hemodynamic responses at this age. In addition, all the studies mentioned above were conducted in high-income countries, limiting generalisability, as responses to paradigms, as well as sleep stage effects, may diverge in low- and middle-income settings due to differences in social and physical environments and possible environmental adaptations.

### 1.1 Variability in hemodynamic responses and relation to sleep stages

Several factors may explain the observed differences in hemodynamic responses between sleep stages. Cerebral blood volume and tissue oxygenation are known to vary across sleep stages, reflecting differences in neural activity and metabolic demand. For example, Doppler flowmetry studies in neonates have shown increased cerebral blood flow during transitions from QS to AS, suggesting elevated metabolic activity associated with AS onset.^34, 35^ Similarly, adult fNIRS studies report elevated oxyhemoglobin concentrations during light sleep and transitions between stages.^36^ However, some neonatal evidence suggests that changes in haemoglobin concentration may not be limited to transition periods: Munger et al., using fNIRS in 2-8-day-old full-term infants, reported fluctuations in oxyhemoglobin concentrations both during and between transitions from AS to QS and vice versa.^37^ Taken together, these findings imply that while some hemodynamic shifts may reflect transitional dynamics, others may be intrinsic to sustained sleep stages themselves.

Supporting this, a longitudinal fNIRS study measuring tissue oxygenation index (TOI), a metric more sensitive to venous saturation and thus oxygen extraction, showed that TOI was lower in AS at 2–4 weeks of age, equivalent across stages at 2–3 months, and greater in AS by 5–6 months.^38^ Given that TOI reflects the balance between cerebral blood flow and oxygen consumption, these findings suggest that AS may initially be characterized by high metabolic demand that outpaces blood flow, leading to greater oxygen extraction. With age, neurovascular regulation matures, and hemodynamic patterns shift to resemble those observed in adults. Thus, the apparent discrepancies across studies may reflect developmental changes in cerebral physiology, rather than true contradictions.

Secondly, sleep stages have been shown to be associated with distinct functional connectivity (FC) patterns in neonates, however, again, findings are mixed.^39–42^ Results of an EEG investigation showed an increase in long-range connectivity in QS, while AS was associated with a local increase in occipital connectivity.^39^ To date, there have been two studies of sleep-stage-dependent changes in FC in newborns using fNIRS. The results of both studies found AS to be characterized by more extensive long-range interhemispheric connectivity, and QS by stronger short-range connectivity.^40, 41^ A recent fMRI investigation of sleep stage effect on functional connectome in the first few weeks of life found no significant difference, although the authors believed their sample to be underpowered and warranted further investigation with a larger sample.^42^

In sum, differences in hemodynamic responses across sleep stages likely reflect both transient and sustained physiological changes, shaped by developmental maturation of neurovascular regulation. Although findings on functional connectivity remain mixed, emerging evidence suggests that QS and AS may support distinct patterns of local and long-range neural communication from early infancy. These physiological and network-level differences raise important questions about how sleep stage might modulate task-evoked brain responses, motivating our investigation of sleep stage effects on response amplitude, stimulus discrimination, and habituation in early development.

### 1.2 Why is it important to examine sleep-stage effect?

Firstly, identifying differences in sensory processing between sleep stages could aid our understanding of the role of AS and QS in functional brain development. Secondly, for stimuli-evoked cortical responses to be a reliable biomarker of pathology or future development, it is important to establish whether the responses and the consistency of measured responses during sleep are sleep stage dependent at each developmental stage. For example, following the discovery of consistent sensory gating in AS but not in QS, Hunter et al. recommended using this response measured only in AS when examining its association with potential risk factors such as nicotine exposure during pregnancy.^19^ Similarly, Nevalainen et al. showed that MEG consistently detects a touch-evoked response in QS, so they recommend that clinicians evaluate somatosensory processing only from data obtained in QS.^9^ When examining group differences, especially when making comparisons between clinical and control groups, it is important to establish that the percentage of recordings in each sleep stage is equivalent across the groups.

### 1.3 Current study

The present study investigates the influence of infant sleep stages on the hemodynamic responses measured with fNIRS during a battery of standard auditory paradigms. The data for this study are part of the Brain Imaging for Global Health (BRIGHT) Project,^43^ a prospective longitudinal study examining infant development from term birth to 24 months of age in the UK and The Gambia (GM). fNIRS data were acquired at each visit at 1, 5, 8, 12, 18, and 24 months of infant age. Only at the one-month visit was fNIRS acquired during natural sleep, and thus only the one-month data are presented here. At both sites, the design included measures of auditory social selectivity, measures of habituation and recovery of response to novelty, and functional connectivity of the resting state. Due to the few infants identified as being in AS compared to QS during functional connectivity analysis (9 AS vs 6 QS in the UK, and 6 AS vs 28 QS in the GM), we did not proceed with comparing connectivity patterns between the two sleep stage groups.

Our first objective was to describe infants’ sleep behavior across the session, and classify them into active sleep and quiet sleep groups for each paradigm. The study was not originally designed to include sleep stage coding, as it was the first longitudinal fNIRS study with infants from sub-Saharan Africa and did not include simultaneous EEG or respiration recordings to reduce the burden on the infants. Instead, we classified sleep stages using detailed behavioral video coding based on established criteria.^5^

Our second objective was to examine how sleep stages modulate hemodynamic responses during two auditory paradigms. After initial scanning of the videos for assessment of video quality, it became apparent that a large number of videos recorded in the GM could not be sleep-stage-classified due to very low lighting used during data collection, which, while optimal for successful fNIRS data collection, obscured most of the classification criteria. This greatly reduced the final sample size. However, the fNIRS data from this time point were initially pre-processed and analyzed by Greenhalgh and colleagues^44^ as part of a larger dataset comprising the full cohort of infants from both sites for a separate investigation into cross-paradigm hemodynamic responses at this age. Therefore, to comprehensively investigate the effects of sleep stages, we employed two complementary analytic approaches: one based on pre-defined regions of interest (ROIs) identified from the full cohort, and a second channel-wise analysis to explore potential sleep-stage differences in the spatial activation patterns and task-evoked responses.

We explored whether sleep stages influenced the amplitude or spatial distribution of these responses, or the strength of non-vocal over vocal discrimination. For the channel-wise analysis of the social selectivity paradigm responses, we expected a higher proportion of infants to exhibit greater hemodynamic response to non-vocal than to vocal stimuli, based on the findings from the full-cohort analysis at one month. Based on prior fNIRS work showing higher response amplitudes in AS than QS regardless of stimulus type,^25^ we hypothesized a generally higher amplitude of hemodynamic response in AS irrespective of condition. However, due to the absence of prior fNIRS studies explicitly testing sleep-stage effects on auditory discrimination, we made no directional hypothesis regarding differences in condition discrimination strength between sleep stage groups.

For the habituation and novelty detection (HaND) paradigm, in line with previous literature^45^ and the results of Greenhalgh et al.,^44^ we expected to see a decrease in response amplitude across repeated presentations of the familiar stimulus irrespective of sleep stage. We assessed sleep stage differences in response amplitude to repeated auditory stimuli, the spatial distribution of these responses, and the degree of habituation. Although no fNIRS studies to date have directly examined sleep stage effects on cortical habituation, EEG literature suggests stronger habituation may occur in AS relative to QS. Thus, we hypothesized a greater habituation in AS.

We did not have specific hypotheses regarding differences between the UK and Gambian cohorts; rather, by examining sleep stage effects in two infant populations, we aimed to strengthen the generalizability of our findings by assessing the consistency of effects across different samples.

## 2 Materials and Methods

### 2.1 Participants

#### 2.1.1 UK cohort

In the UK, 62 families were recruited at the Rosie Hospital, Cambridge University Hospitals NHS Foundation Trust at 32 - 36 weeks’ gestation, with one family withdrawing before the one-month time point. Once per week during the recruitment phase, all families visiting for their antenatal visit with a healthy pregnancy were approached and given information about the study. Interested families were followed up via email/phone and recruited into the study. Inclusion criteria included healthy, full-term pregnancy (37-42 weeks gestation), birth weight *>* 2.5*kg* and absence of any known genetic or neurological conditions. The majority of families lived either in the university town or in surrounding urban or rural communities within a 20-mile radius. The study was approved by the National Research Ethics Service East of England Committee, NHS Health Research Authority (REC reference 13/EE/0200), and informed written consent was obtained from all parents before participation.

#### 2.1.2 Gambian cohort

Detailed characteristics of the Gambian site, context, and recruitment strategies have been described in detail previously.^43^ Briefly, the Gambia, located on the West coast of Africa and bordering Senegal, is largely rural, with many families practicing subsistence farming and living in extended households. Despite historically low national income and education levels, school attendance has improved significantly in recent decades. Childcare is typically shared among family members, and Islam plays a central role in shaping family and community life. The Gambian arm of the BRIGHT project was conducted at the Keneba Field Station of the Medical Research Council The Gambia (MRCG) at the London School of Hygiene and Tropical Medicine (LSHTM) in rural West Kiang, a region affected by pronounced seasonal variation in food availability. Participants were recruited from the rural village of Keneba and nearby villages within a 20 km radius. Over the study period, local infrastructure saw notable improvements, such as new roads, enhancing connectivity for the previously more isolated field station. Pregnant women were identified using the West Kiang Demographic Surveillance System and invited to an antenatal clinic visit to assess eligibility if they expressed interest in taking part in the study. Eligibility criteria required women to have a singleton pregnancy, with Mandinka as their primary language, and to be less than 36 weeks pregnant by the time of the first antenatal visit. Infants were included if born at term (37–42 weeks’ gestation). In total, 222 families were recruited into the study at delivery, with 204 families still enrolled at the time of the one-month visit (as 8 were stillborn, 7 suffered neonatal death and 3 withdrew post birth - see the protocol paper for detail^43^). Ethical approval was granted by the joint Gambia Government – MRC Unit and the Gambia Ethics Committee and the Scientific Coordinating Committee at the MRC Unit The Gambia (SCC 1351). Informed consent was obtained from all parents before participation in writing or via thumbprint.

### 2.2 Experimental procedure and stimuli

The BRIGHT study one-month session comprised of a battery of neuroimaging (EEG, fNIRS), behavioral (Neonatal Behavioral Assessment Scale (NBAS), parent-child interaction), and anthropometric measures. The analyses reported in the current work focus on the fNIRS data. The infants were allowed to naturally fall asleep at some point during their visit. A researcher would wrap the fNIRS headband around the infant’s head before or after the infant fell asleep. During the presentation of the fNIRS paradigms, infants were held by a researcher or parent seated on a comfortable chair. If an infant began to wake, the researcher or parent would attempt to rock them back to sleep. Aside from this soothing, they were instructed not to interact with the infant. At both data collection sites, testing room lights were dimmed or switched off to aid infants’ sleep and comfort. Sessions were video recorded using a camera on a tripod positioned approximately 30 cm from the infant, with at least the head and torso in view, to monitor behavior and sleep. The fNIRS recording always followed the same order of auditory paradigms: social selectivity, followed by habituation and novelty detection (HaND). Stimuli were presented using a MATLAB® custom-written stimulus presentation framework, Task Engine,^46^ and Psych toolbox on an Apple Macintosh computer. The intensity of the sound presented via Logitech Z130 stereo speakers was adjusted to a measured ∼ 60 dB at the position of the infant’s head (range across the task: 60.1 dB – 61.4 dB).

### 2.2.1 The social selectivity paradigm

The sequence of stimulus presentation has been used in previous research.^47–49^ Briefly, two types of conditions (*vocal (V)*, *non-vocal (N))*, were presented for eight trials each, with a 10-12 second silent baseline period in-between. The two types of conditions were presented in the same pseudo-random order across infants and were identical across the UK and the GM sites. The vocal condition comprised four non-speech adult vocalizations of two speakers (who coughed, yawned, laughed, and cried). The non-vocal condition comprised four common environmental sounds (running water, toys, bells and rattles). Both conditions were 8 seconds long.

#### 2.2.2 The habituation and novelty detection paradigm

The HaND stimuli has been previously described in Lloyd-Fox et al.^45^ Briefly, they consisted of spoken sentences (UK: ‘“Hi baby! How are you? Are you having fun? Thank you for coming to see us today. We’re very happy to see you’, Gambian: “Denano a be nyadii. I be kongtan-rin? Abaraka bake ela naa kanan njibee bee, n kontanta bake le ke jeh”). Two versions of the stimulus were recorded in each language by native English or Mandinka male and female speakers. Each sentence was considered one trial, 8 seconds long, preceded by a 10-second silent baseline, given in a blocked design. The trials were presented in the following order: 15 repetitions of a female speaker; 5 repetitions of a male speaker; and 5 repetitions of the same female speaker. These trials were clustered into the following epochs: familiarization rials 1 – 5 (Fam1); familiarization trials 6 – 10 (Fam2); familiarization trials 11 – 15 (Fam3); novel trials 16 – 20 (Novel) and trials 21 – 25 (Post-test).

### 2.3 fNIRS data acquisition

fNIRS data were collected using a continuous wave NTS fNIRS system (Gowerlabs Ltd., London, UK^50^). The system makes use of two wavelengths of near-infrared light (780 and 850 nm) to detect changes in oxyhemoglobin (HbO) and deoxyhemoglobin (HbR) concentrations, using a sampling frequency of 10 Hz. Infants were assessed using a custom-built fNIRS headgear covering both hemispheres and consisting of an array of 8 sources and 8 detectors, for a total of 18 channels (9 per hemisphere), covering bilateral frontal and temporal regions, with a source-detector separation of 2 cm (Fig.1). For each infant, head measurements were taken to enable the alignment of the headgear with the placement coordinates.^51^ Photographs of the participants were taken after the headbands were secured on the participant’s head, and again at the end of the session to facilitate

**Fig 1.**
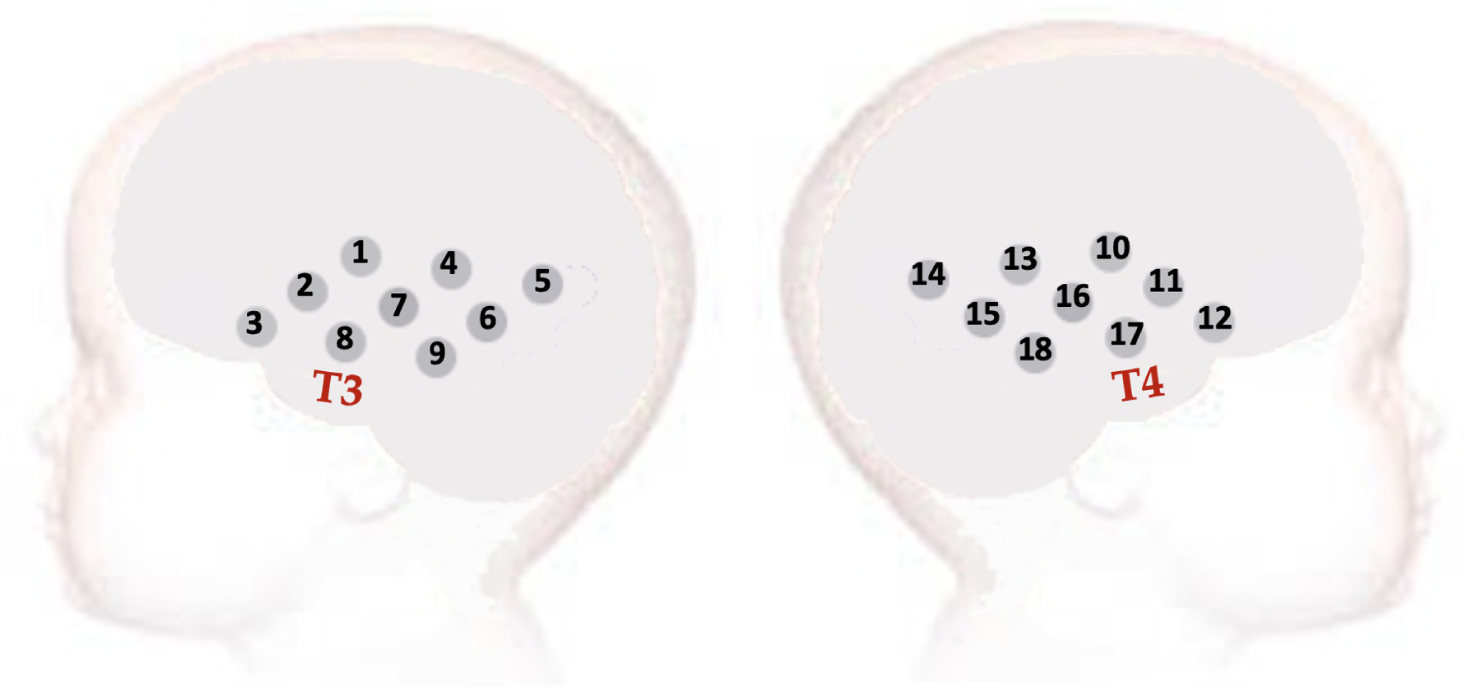
The 18-channel fNIRS array used at the 1-month visit, covering bilateral frontal and temporal regions, with the 10-20 system T3 and T4 landmarks clearly marked offline coding of the headgear placement and identification of measurement location relative to external anatomical landmarks. With this information, the underlying cortical anatomy of the fNIRS channels was approximated in this study.

### 2.4 fNIRS preprocessing

The data used in the current analysis was pre-processed by Greenhalgh and colleagues^44^ as part of a separate investigation using the full dataset. Individual channel level quality was assessed using QT-NIRS function,^52^ with filtering parameters adjusted for infant heart rates (cut-off frequencies: [1.5, 3.5] Hz, SCI: 0.7, PSP: 0.1 µV). Datasets with more than 40% low-quality channels (based on SCI/PSP thresholds) were excluded. Remaining data were converted from intensity to optical density, then underwent motion detection by channel using Homer2^53^ function (using thresholds: tMask: 1, tMotion: 1 s, STDEVthresh: 15, AMPthresh: 0.5) and trial rejection if *>* 50% of channels showed motion. Motion correction used spline interpolation (*p* = 0.99) and wavelet filtering (*IQR* = 0.8). Data were then low-pass filtered (0.6*Hz*) and converted to changes in HbO and HbR concentrations using the modified Beer-Lambert Law (DPF: 5.22, 4.23). Block averaging and linear detrending were applied (tRange: -4 to 20 s). Only datasets with at least three valid trials per condition (i.e. 3 N and 3 V trials in the social selectivity paradigm, and 3 Fam1, 3 Fam3 and 3 Novel trials in the HaND paradigm) were retained. The HbO and HbR values for each channel were extracted for the sub-sample of infants with valid sleep stage coding and used for the analyses presented here.

### 2.5 Sleep stage coding

Although the terms *sleep state* and *sleep stage* are often used interchangeably in studies of infant sleep, here we adopt the term *sleep stage* to refer specifically to active sleep (AS) and quiet sleep (QS). This distinction aligns with developmental sleep physiology literature, which classifies AS and QS as distinct stages of infant sleep architecture, each characterized by unique behavioral, cardiovascular, and neural patterns. Throughout this paper, we therefore use *sleep stage* to denote these two neurophysiologically and functionally discrete phases.

Video recordings of infants during the fNIRS paradigms were reviewed and micro-coded offline to classify periods during which each infant was in AS, QS, or awake. The video was coded in segments of 15 seconds, following the infant states classification described in the Neonatal Behavioral Assessment Scale,^5^ which defines a state as being achieved if the infant remains in that state for at least 15 seconds. Prior to classifying the infant’s sleep stages, the coders watched two minutes of video recording directly preceding the start of each paradigm to familiarize themselves with the infant’s behavior.

Quiet sleep was scored if the infant’s eyes were closed without rapid eye movements and met at least one additional criterion: regular respiration, no spontaneous activity except (rapid suppression of) startles or movements at regular intervals. Active sleep was scored if the infant’s eyes were closed, though possibly opening briefly, with at least one additional criterion: rapid eye movements (REMs), low (muscle) activity with random movements and startles or startle equivalents, smoother movements compared to QS, irregular respiration, or intermittent sucking movements. If the infant was not in either sleep stage (e.g., fussy or calmly awake), the state was coded as awake. Each criterion was assigned a confidence level based on how important it was shown to be for sleep stage discrimination in newborns, based on previous literature.^54^ If a behavior could not be coded due to factors such as poor visibility, occlusion by blankets, or low lighting, the confidence level for that segment was reduced. Behaviors that were mutually exclusive between sleep stages (e.g., REM vs. non-REM, regular vs. irregular respiration) contributed more strongly to the confidence rating. When such key features were unobservable (e.g., due to infant position or obstruction), coders’ confidence in classifying the sleep stage for that epoch was substantially reduced. Conversely, behaviors that may occur in both sleep stages (e.g., startles) contributed less to confidence, and their absence, such as when the infant’s arms were held, did not significantly lower confidence if other informative behaviors were visible. Confidence scores were calculated for each 15-second segment, ranging from 0% to 100% (Supplemental Tab. 1). Infants with an average confidence score below 65% were excluded from the analyses. Sequentially, infants that remained in one stage throughout the paradigm were allocated to either AS or QS group. If the infant transitioned between stages during the paradigm, they were placed in a third ’transition’ group.

Data from infants in transition group were inspected for whether they had at least 3 valid consecutive trials of each condition with continuous sleep stage. Firstly, since the infant’s state was coded in 15-second segments, it had to be aligned with the timing of the fNIRS trials, which were 18-20 seconds long in the social selectivity paradigm and 18 seconds long in the HaND paradigm. Behavioral coding was aligned with the timings of the fNIRS paradigm by marking the start of each trial when the auditory stimulus was first heard. For both the social selectivity and the HaND paradigms, if an infant transitioned within the trial itself, the trials were labeled with the stage that occupied the longest duration within that trial. In the case of the social selectivity paradigm, if an infant had at least three valid N and three valid V trials within one continuous block of 6 trials in one sleep stage, these trials were retained for the analysis while the trials with another sleep stage were excluded. In the case of the HaND paradigm, if an infant remained in the same sleep stage during the familiarization trials with at least three valid Fam1 and three valid Fam3 trials in this sleep stage, they were retained for the habituation analysis. Similarly, if infants had at least three valid Fam3 trials directly followed by at least three valid Novel trials in the same sleep stage, they were retained for the novelty detection analysis.

### 2.6 Statistical analyses

All statistical analyses were performed in R.^55^ When assumptions of any test were violated, a non-parametric alternative was performed. Benjamini-Hochberg correction^56^ was used to control the False Discovery Rate (FDR) (*α* = 0.05). Consistent with prior infant fNIRS research,^48^ we treated a significant increase in HbO and/or a significant decrease in HbR as an indicator of cortical activation. If, however, both chromophores showed significant changes in the same direction - either increasing or decreasing together - this was considered atypical for a functional hemodynamic response and excluded from interpretation. To compare the quality of the fNIRS data across sleep stages we computed two metrics: 1. percentage of clean data - defined as the percentage of data out of the total length of the recording (per paradigm) that did not require motion correction; 2. number of valid trials retained per condition. Independent samples t-tests were used to compare these metrics between AS and QS groups within each site.

Two complementary analytical approaches were employed to examine the influence of sleep stages on infants’ hemodynamic responses within the two paradigms. The first, referred to as the region of interest (ROI)-based approach, focused on predefined sets of channels that showed significant activation and condition contrasts for each paradigm separately, based on analyses of the full cohort of infants across both sites, as reported by Greenhalgh et al.^44^ This approach did not account for sleep stage in the selection of channels but benefited from the increased statistical power of the larger sample. In Greenhalgh et al.,^44^ significant channels were identified using threshold-free cluster enhancement (TFCE; *E* = 0.5, *H* = 2) - a method that addresses multiple comparison issues and enables detection of significant activation patterns without requiring apriori cluster definitions.^57^ The time windows and ROIs derived from the TFCE analysis were then applied to the subsample of infants with valid sleep stage coding to investigate sleep stage-related differences using this ROI-based approach.

The second, referred to as the channel-wise analysis, involved an exploratory investigation of sleep stage effects across all channels within the smaller subset of infants for whom sleep stages were reliably coded. This approach enabled assessment of whether hemodynamic responses varied by sleep stage, albeit with reduced statistical power due to the smaller sample size.

#### 2.6.1 Effect of sleep stages on the amplitude of significant activations with the ROI-based approach

The ROI-based approach was first applied to assess the effect of sleep stage on the amplitude of significant activations in the social selectivity and HaND paradigms.

In Greenhalgh et al.,^44^ for social selectivity, channels with a significant activation to N and V trials compared to baseline were identified across the full UK (N = 46) and Gambian (N = 148) cohorts. To maintain comparable signal-to-noise ratios across conditions, only channels that exhibited significant activation to both N and V stimuli were included, ensuring an equal number of channels for each condition. Hence, for the UK cohort, the response to each condition was averaged across HbO channels 9, 15, 16, 18, and HbR channels 15 and 18 within each sleep stage group. For the GM cohort, the response to each condition was averaged across HbO channels 15, 16, 18, and HbR channels 6, 7, 9, 10, 13, 15, 16, 18 in each sleep stage group. Independent samples t-tests were conducted for each site separately, comparing AS and QS groups within each social selectivity condition.

To investigate the effect of sleep stage on the amplitude of HbO and HbR responses in the HaND paradigm, we focused on the average responses during Fam1, before habituation could take place. Therefore, we used channels identified as having significant activation to Fam1 compared to baseline in Greenhalgh et al across the full UK (N = 38) and GM (N = 136) cohorts. For the UK cohort, Fam1 response was averaged across HbO channels 4 and 7 and HbR channels 7, 15, 18 in each sleep stage group. For the GM cohort, Fam1 response was averaged across HbO channels 1, 4, 7, 13 and HbR channels 4, 7, 13 in each sleep stage group. Independent samples t-tests were conducted for each site separately, comparing average Fam1 hemodynamic response between AS and QS groups.

#### 2.6.2 Effect of sleep stages on the strength of the responses for contrasts of interest with the ROI-based approach

Next, the ROI-based approach was applied to assess whether sleep stage influenced the strength of non-vocal selectivity - that is, the degree to which infants responded more strongly to non-vocal than vocal stimuli. The significant regions identified in Greenhalgh et al.^44^ in the full UK cohort for the N *>* V selectivity were over bilateral posterior temporal regions of the array. Concretely, significant N *>* V selectivity was present in five HbO channels (channels 4, 13, 14, 15, 16) and nine HbR channels (channels 4, 7, 9, 13, 14, 15, 16, 17, 18). There were no channels with significant V *>* N selectivity. For the Gambian cohort, no N *>* V or V *>* N selective regions were observed in the full cohort. Therefore, for the current work, ROI-based analyses of the effect of sleep stage on the strength of N *>* V selectivity were focused on the UK cohort. After averaging the HbO and HbR responses across the selected channels, condition contrast values were computed for each infant by subtracting the mean V response from the mean N response. These contrast values were then compared between AS and QS groups using an independent samples t-test.

Similarly, the ROI-based approach was applied to assess whether sleep stage influenced the degree of habituation in the HaND paradigm. The regions with significant habituation (Fam1 *>* Fam3) identified in Greenhalgh et al.^44^ in the full UK cohort were observed over posterior parts of the array. Significant habituation was present in two HbO channels (channels 4 and 7) and one HbR channel (channel 7). A response to novel stimulus after the habituation phase (Novel *>* Fam3) was not present. For the Gambian cohort, the significant regions were observed in more central temporal regions of the array. Significant habituation was present in two HbO channels (channels 1 and 7) and three HbR channels (channels 4, 7 and 13). As with the UK cohort, no Novelty response at group level was observed. Therefore, ROI-based investigation of the sleep stage effect focused on the Fam1 *>* Fam3 response in both cohorts. We computed the contrast value for each infant by subtracting the mean Fam3 response from the mean Fam1 response. These contrast values were then compared between the AS and QS groups using an independent sample t-test.

#### 2.6.3 Effect of sleep stages on the fNIRS responses with the channel-wise approach

For the channel-wise analyses, the hemodynamic responses within the QS and AS infants were assessed separately. First, we identified the channels with significant activation to each condition compared to baseline using one-sample t-tests. Next, the channels that showed a significant response to condition (N *>* baseline, Fam1 *>* baseline, Novel *>* baseline) were examined for selectivity to paradigm conditions (N *>* V in the social selectivity paradigm, or Fam1 *>* Fam3 and Novel *>* Fam3 for the HaND paradigm), using paired-sample t-tests.

## 3 Results

### 3.1 Characteristics of the UK and Gambian cohorts

Only participants who contributed valid fNIRS and sleep stage data were retained for the current analysis. The reasons for participant retention and exclusion are outlined below.

#### Gambian cohort

Of the total number of participants eligible at 1 month (*N* = 204), 136 had valid fNIRS data for at least the habituation part (Fam1, Fam2, Fam3) of the HaND paradigm and 148 had valid fNIRS data for the social selectivity paradigm. Participant retention per paradigm based on fNIRS data quality in the two cohorts is presented in Supplemental Fig. 1. For the social selectivity paradigm, a total of 46 infants contributed both valid fNIRS and sleep stage data (Fig. 2). Seven infants (15%) transitioned between sleep stages, 16 infants (35%) remained in QS and 23 infants (50%) in AS during the entire task. The infants that changed between sleep stages were retained for further analysis if they had at least three trials of each condition (N and V) in the same sleep stage. If an infant changed sleep stage during the paradigm, their data from only one sleep stage were used. In other words, they could not have contributed data to both AS and QS groups, to avoid the potential confounding that a process of changing state could bring on the hemodynamic responses to the paradigm. The resulting sample included 20 QS and 26 AS infants. For the HaND paradigm, a total of 52 infants contributed both fNIRS and sleep stage data (Fig. 2). Eleven infants (21%) transitioned between sleep stages, 25 infants (48%) remained in QS and 16 infants (31%) remained in AS throughout the entire task (Fam1 - Novel trials). The infants that transitioned between sleep stages during the paradigm were retained for habituation (Fam1 - Fam3) or novelty detection (Fam3 - Novel) analyses if they had at least six consecutive trials in one sleep stage across the two epochs (at least three trials in each epoch). Hence, there were 27 QS and 24 AS infants that contributed data for the analysis of Fam1 response, 25 QS and 20 AS infants that contributed the data for the habituation (Fam1 to Fam3) analysis, and 29 QS and 18 AS that contributed the data for the novelty detection analysis (Fam3 to Novel). The number, age, and sex of participants who contributed data for the analysis are presented in Table 1.

**Fig 2.**
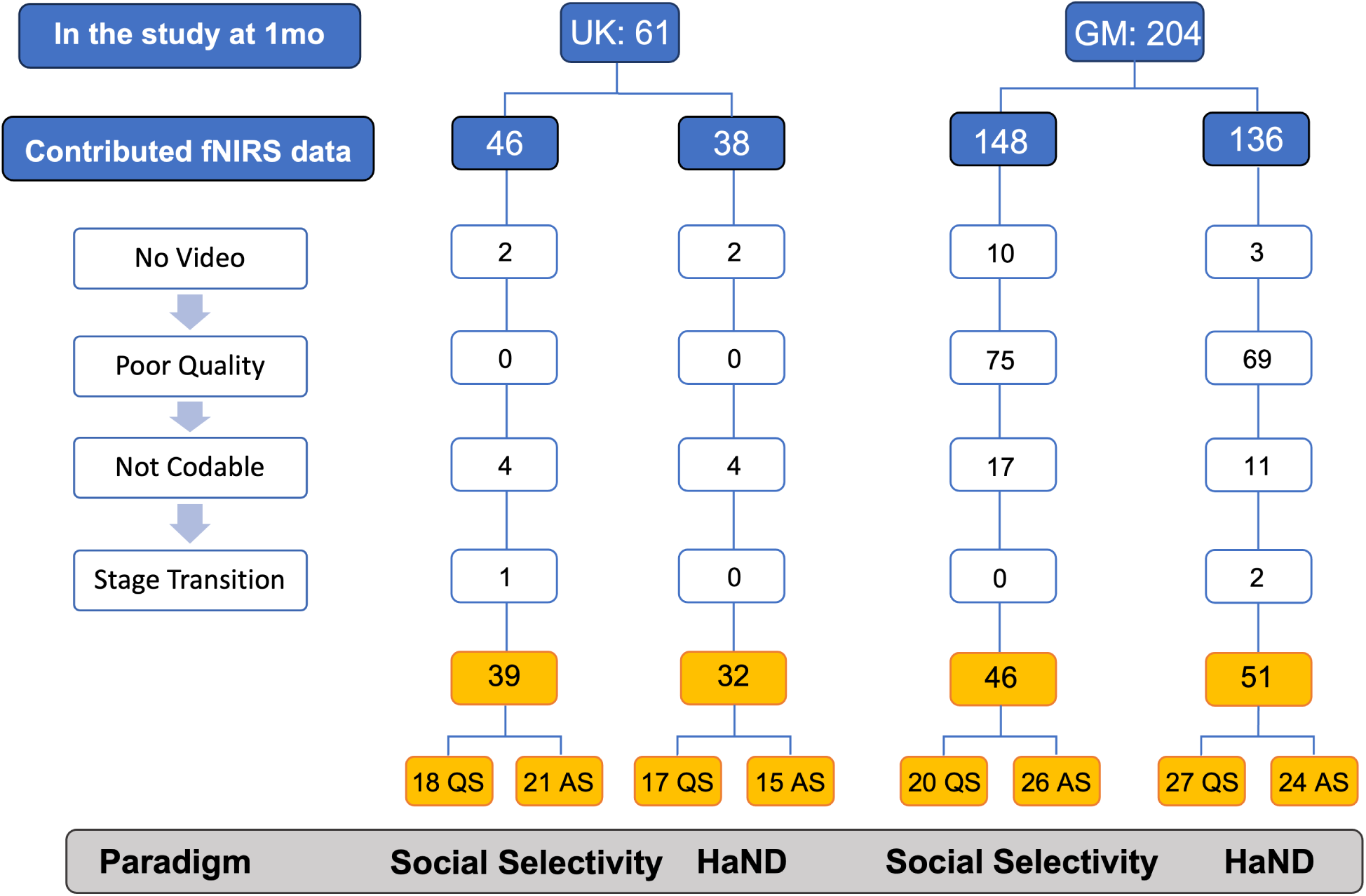
Reasons for exclusion of participants from the United Kingdom (UK) and Gambian (GM) cohorts from the final analysis across the two paradigms: social selectivity and Habituation and Novelty Detection (HaND). The reasons for exclusion could be: No Video - video of the whole session or the specific task is missing; Poor Quality - behaviors cannot be coded because of insufficient lighting in the testing room; Not Codable - behaviors cannot be coded from the video because infant is facing away from the camera, is wrapped up or in the position that makes coding behaviors not possible; Stage Transition - infant transitioned from one sleep stage to another during the paradigm and there are not enough trials of each condition (for social selectivity and HaND) in either stage.

**Table 1.** Number of UK and Gambian (GM) infants, their mean age (days) and gender (M = male, F = Female) that contributed fNIRS and sleep stage data for each paradigm.

| Site | paradigm | Sleep Stage | N | Mean Age (SD), days | Sex |
| --- | --- | --- | --- | --- | --- |
| UK | Social selectivity | QS | 18 | 31.6 (5.2) | 8 F, 10 M |
|  |  | AS | 22 | 34.3 (6.7) | 10 F, 12 M |
|  | HaND | QS | 16 | 33.7 (7.3) | 6 F, 10 M |
|  |  | AS | 15 | 32.4 (5.4) | 10 F, 5 M |
| GM | Social selectivity | QS | 20 | 35.1 (4.9) | 10 F, 10 M |
|  |  | AS | 26 | 37.0 (6.9) | 14 F, 12 M |
|  | HaND | QS | 25 | 36.4 (6.0) | 14 F, 11 M |
|  |  | AS | 20 | 37.1 (6.0) | 11 F, 9 M |

**Table 2.**
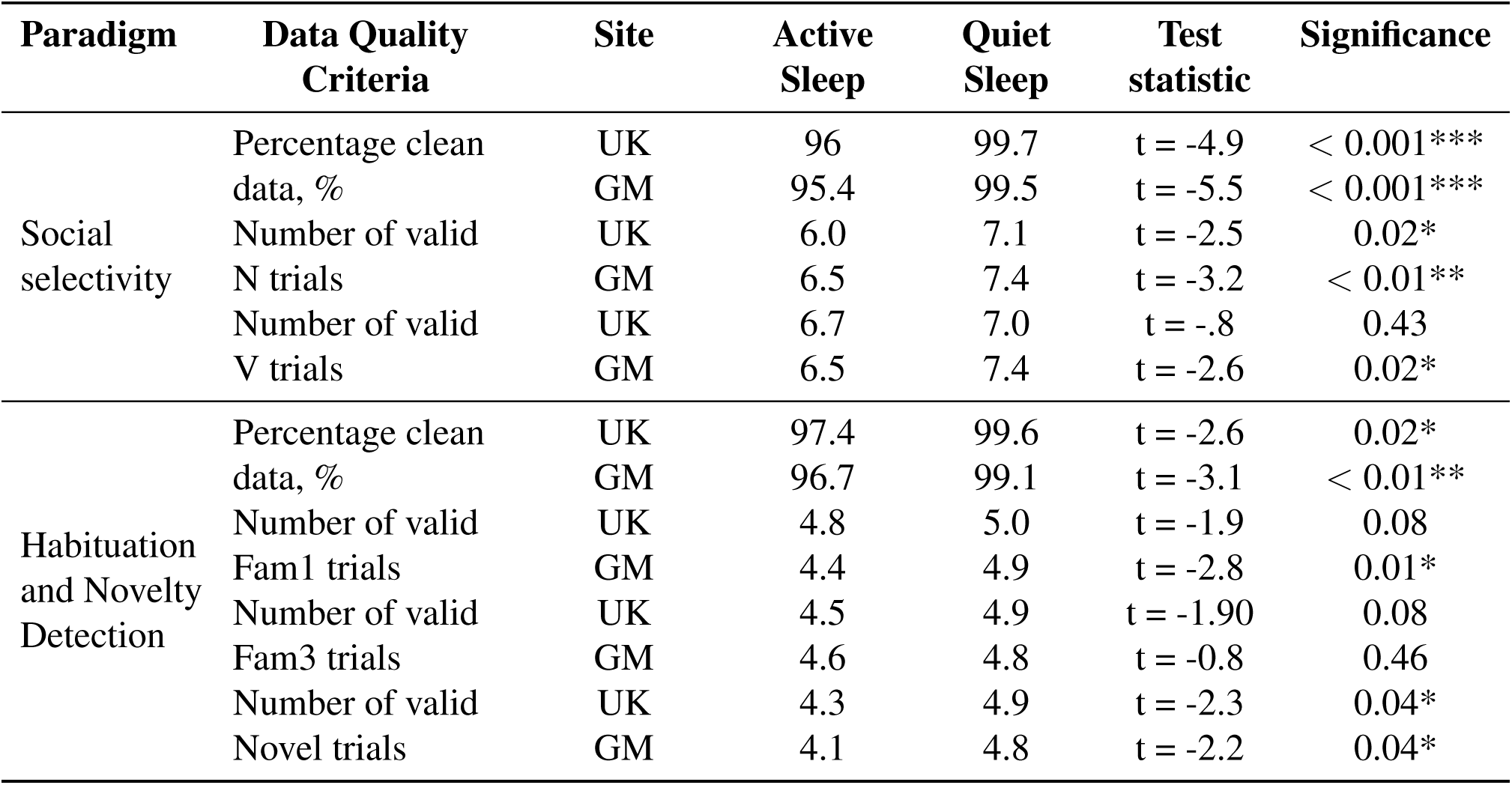
Data quality comparison according to sleep stage for each paradigm in the UK and Gambian (GM) cohorts.

| Paradigm | Data Quality Criteria | Site | Active Sleep | Quiet Sleep | Test statistic | Significance |
| --- | --- | --- | --- | --- | --- | --- |
| Social selectivity | Percentage clean data, % | UK | 96 | 99.7 | $t = -4.9$ | $< 0.001^{***}$ |
| | | GM | 95.4 | 99.5 | $t = -5.5$ | $< 0.001^{***}$ |
| | Number of valid N trials | UK | 6.0 | 7.1 | $t = -2.5$ | $0.02^*$ |
| | | GM | 6.5 | 7.4 | $t = -3.2$ | $< 0.01^{**}$ |
| | Number of valid V trials | UK | 6.7 | 7.0 | $t = -.8$ | $0.43$ |
| | | GM | 6.5 | 7.4 | $t = -2.6$ | $0.02^*$ |
| Habituation and Novelty Detection | Percentage clean data, % | UK | 97.4 | 99.6 | $t = -2.6$ | $0.02^*$ |
| | | GM | 96.7 | 99.1 | $t = -3.1$ | $< 0.01^{**}$ |
| | Number of valid Fam1 trials | UK | 4.8 | 5.0 | $t = -1.9$ | $0.08$ |
| | | GM | 4.4 | 4.9 | $t = -2.8$ | $0.01^*$ |
| | Number of valid Fam3 trials | UK | 4.5 | 4.9 | $t = -1.90$ | $0.08$ |
| | | GM | 4.6 | 4.8 | $t = -0.8$ | $0.46$ |
| | Number of valid Novel trials | UK | 4.3 | 4.9 | $t = -2.3$ | $0.04^*$ |
| | | GM | 4.1 | 4.8 | $t = -2.2$ | $0.04^*$ |

#### UK cohort

Of the total participants eligible at 1 month (*N* = 61), 46 had valid fNIRS data for the social selectivity paradigm and 38 for the HaND paradigm (Supplemental Fig. 1). During the social selectivity paradigm, out of 40 infants with valid sleep stage classification (Fig.2), 8 infants (20%) transitioned between sleep stages, 14 infants (35%) remained in QS and 18 (45%) in AS during the entire paradigm. After incorporating valid trials from infants that changed between sleep stages, the final sample comprised 18 QS and 21 AS infants. During the habituation paradigm, out of 32 infants with valid sleep stage classification (Fig. 2), 3 infants (9%) transitioned between sleep stages, 13 infants (41%) were in AS and 16 (50%) in QS. All three infants that changed stage were retained for further analysis, bringing the total sample size to 32 infants (17 QS and 15 AS) for the analysis of Fam1 response, 16 QS and 15 AS infants contributed data for the habituation (Fam1 to Fam3) analysis, and to 16 QS and 14 AS were retained for novelty detection (Fam3 to Novel) analysis. The number, age, and sex of participants who contributed data for the analysis are presented in Table 1.

### 3.2 Data quality according to sleep stage

Data quality was sufficiently high for both paradigms in both sleep stage groups (*>* 95%) (Tab.2). Statistical comparison tests showed that for AS, the data quality was significantly lower compared to QS. The number of valid trials retained for analysis was also significantly lower in AS compared to QS for both social selectivity paradigm conditions (N and V) in both cohorts, except the number of valid V trials in the UK. For the HaND paradigm, significantly more valid Fam1, Fam3 and Novel trials were retained for QS compared to AS infants in the GM cohort. In the UK cohort, a significant difference was found only for the Novel trials with more valid trials in QS than AS group. Nevertheless, all data quality ranged between 95 and 100 percent, indicating that the data quality was still very high in both sleep stage groups.

### 3.3 Results of the ROI-based analyses

#### 3.3.1 Sleep stage effect on the amplitude of significant responses

We compared the mean amplitude of the hemodynamic response to vocal and non-vocal conditions of the social selectivity paradigm, and to the first familiarization epoch of the HaND paradigm between sleep stage groups. For the social selectivity paradigm, we found no significant differences in the mean response amplitude by sleep stage for any of the conditions in the UK or GM cohorts for HbO (Fig. 3) or HbR (Supplemental Fig. 2) (all *p >* 0.46).

**Fig 3.**
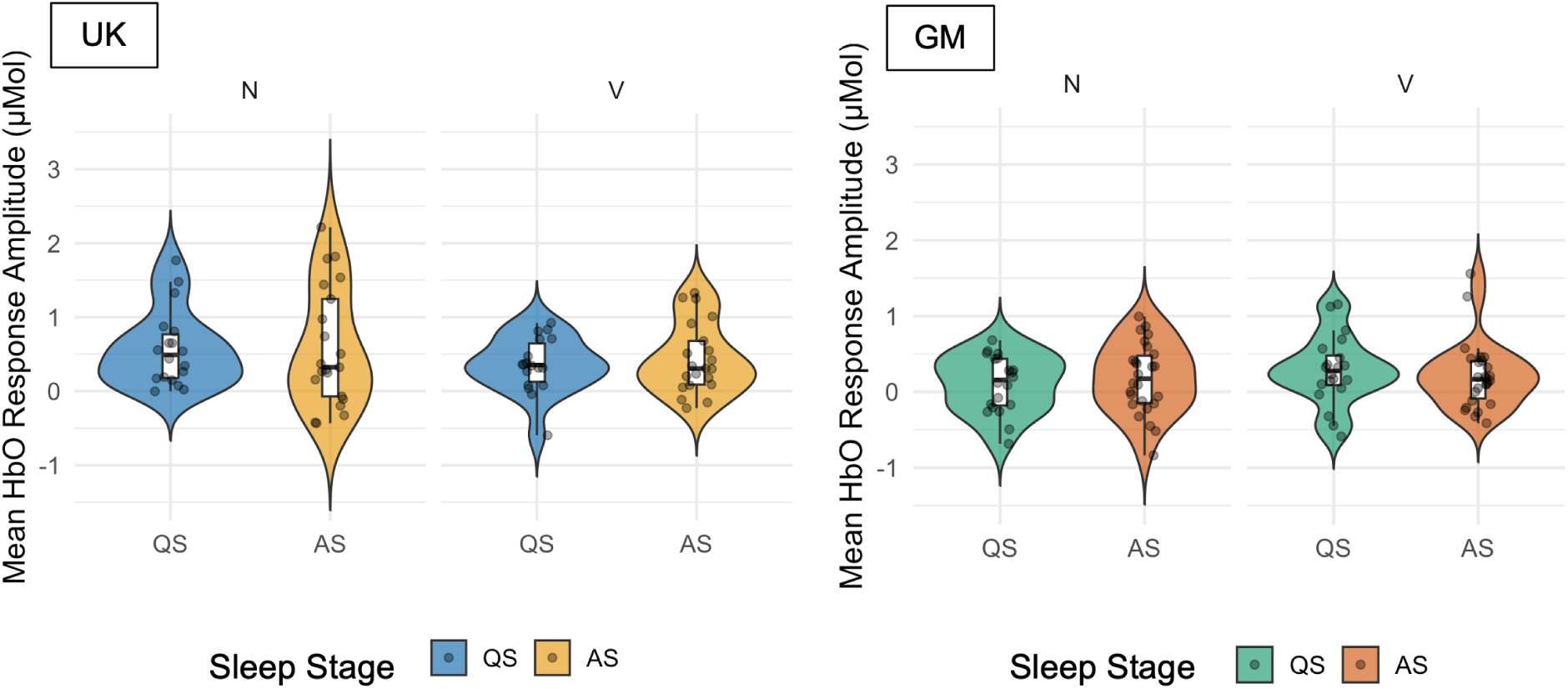
The distribution of mean HbO response amplitudes (*µ*Mol) to non-vocal (N) and vocal (V) trials of the social selectivity paradigm across sleep stages (QS = Quiet Sleep, AS = Active Sleep) in the UK (left) and Gambian (GM) (right) cohorts. Each plot illustrates individual data points, overall distributions, and embedded boxplots.

For the HaND paradigm in the UK cohort, the HbO responses during Fam1 were found to be significantly higher in AS compared to QS (QS *M* = 0.14*µ*Mol*, SD* = 0.39; AS *M* = 0.57*µ*Mol*, SD* = 0.46*, t*(28) = −2.79*, p* = 0.01) (Fig. 4). The effect was not present in HbR (*p* = 0.51) (Supplemental Fig. 3). In the GM cohort, we did not observe an effect of sleep stage in the HbO or HbR responses during Fam1 (all *p >* 0.8) (Fig. 4 and Supplemental Fig. 3)

**Fig 4.**
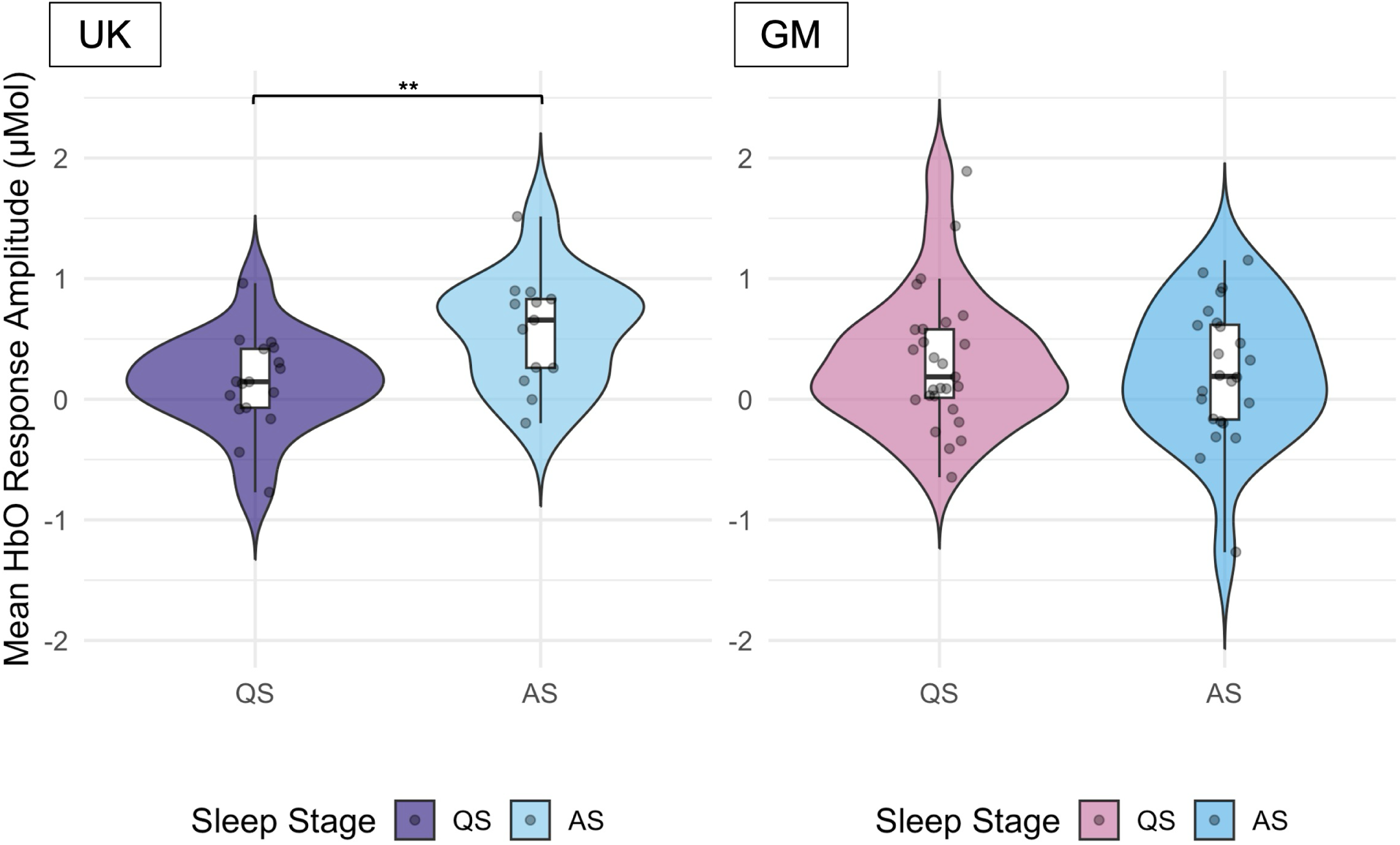
The distribution of mean oxyhemoglobin (HbO) response amplitudes (*µ*Mol) to first five familiarization trials (Fam1) of the habituation and novelty detection (HaND) paradigm across sleep stages (QS = Quiet Sleep, AS = Active Sleep) in the UK (right) and Gambian (GM) (left) cohorts.

#### 3.3.2 The effect of sleep stages on the strength of non-vocal selectivity

A two-sample t-test comparing average N–V condition contrast values between sleep stage groups revealed no significant difference for HbO (QS *M* = 0.32*µ*Mol*, SD* = 0.52; AS *M* = 0.25*µ*Mol*, SD* = 0.86*, t*(37) = 0.27*, p* = 0.79, *Cohen’s d* = 0.09 (negligible), 95% CI [-0.41, 0.53]) or HbR (QS *M* = −0.11*µ*Mol*, SD* = 0.13; AS *M* = −0.12*µ*Mol*, SD* = 0.19*, t*(37) = 0.25*, p* = 0.80, Cohen’s d = 0.08 (negligible), 95% CI [-0.11, 0.14]) in the UK cohort (Supplemental Fig. 4).

#### 3.3.3 The effect of sleep stages on the habituation strength

In the UK cohort, a two-sample t-test revealed a significant HbO Fam1 - Fam3 difference across sleep stages (QS *M* = −0.12*µ*Mol*, SD* = 0.51; AS *M* = 0.49*µ*Mol*, SD* = 0.58, *t*(25) = −2.92*, p* = 0.007, *Cohen’s d* = −1.1 (large), 95% CI [-1.04 -0.18]) (Fig. 5). The difference was not significant for HbR (QS *M* = −0.04*µ*Mol*, SD* = 0.22; AS *M* = −0.20*µ*Mol*, SD* = 0.38*, t*(28) = 1.43*, p* = 0.16, *Cohen’s d* = 0.52 (moderate), 95% CI [-0.07 0.39]) (Supplemental Fig. 5).

**Fig 5.**
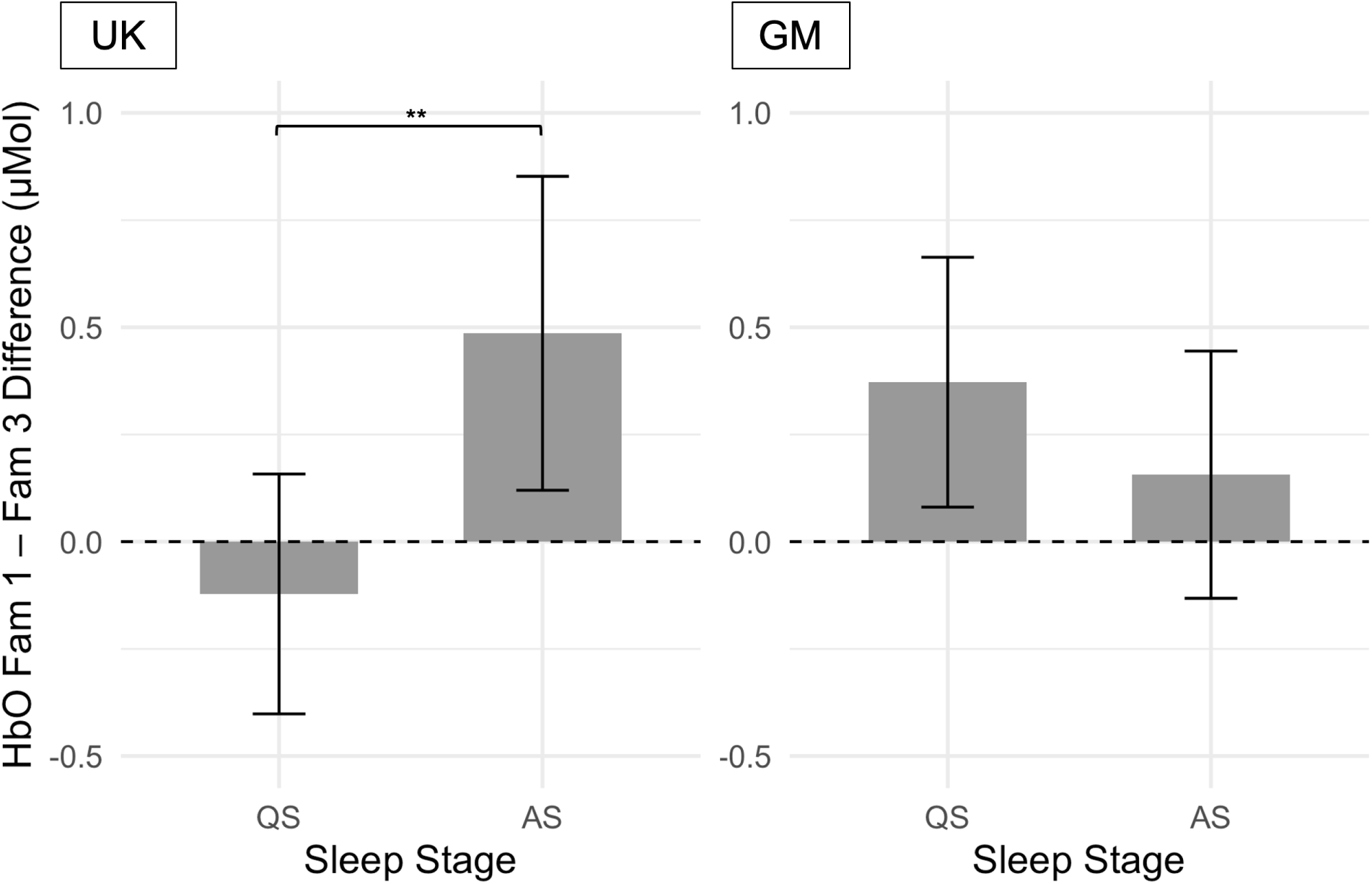
Bar plot showing the mean difference (Fam1 – Fam3) in oxyhemoglobin (HbO) concentration for quiet (QS) and active (AS) sleep groups in the UK (right) and Gambian (GM) (left) cohorts. Error bars indicate standard errors.

In the GM cohort, there were no significant differences in habituation values between sleep stages for HbO (QS *M* = 0.37*µ*Mol*, SD* = 0.71; AS *M* = 0.16*µ*Mol*, SD* = 0.61*, t*(43) = 1.08*, p* = 0.29, *Cohen’s d* = 0.32 (small), 95% CI [-0.19 0.62]) (Fig. 5) or HbR (QS *M* = −0.12*µ*Mol*, SD* = 0.22; AS *M* = −0.06*µ*Mol*, SD* = 0.19*, t*(43) = −0.94*, p* = 0.35, *Cohen’s d* = −0.28 (small), 95% CI [-0.19 0.07]) (Supplemental Fig. 5).

### 3.4 Channel-wise analyses

#### 3.4.1 Sleep-stage specific spatial distribution of significant response to the social selectivity paradigm

First, we determined the significance of the auditory response to V and N conditions separately in the AS and QS groups (one-sample t-test, signal *>* 0, FDR-corrected for the entire set of channels, *p adj <* 0.05) (Fig. 6). In the UK cohort, in the QS group, significant activation was found in 6 out of 18 HbO channels (∼33%) and 3 HbR channels (∼17%) to V condition; activation to N condition was found in 10 HbO channels (∼56%) and 14 HbR channels (∼78%). In the AS group, significant activation to V condition was found in 4 HbO channels (∼22%) and 0 HbR channels; activation to N condition was found in 7 HbO channels (∼39%) and 5 HbR channels (∼28%). A significant selectivity towards N *>* V condition (paired-sample t-test, two-tailed, FDR-corrected for the set of channels significant against baseline, *p adj <* 0.05) was detected in 4 HbR channels (∼22%) in QS (5, 9, 13, 14) and 3 HbR channels (∼17%) in AS (4, 5, 16), but not in any HbO channels (Table 3). There were no channels with significant V *>* N selectivity. In the GM cohort, no channels showed significant HbO or HbR activation to either vocal or non-vocal condition, hence N *>* V selectivity was not investigated in our GM sub-sample of infants.

**Fig 6.**
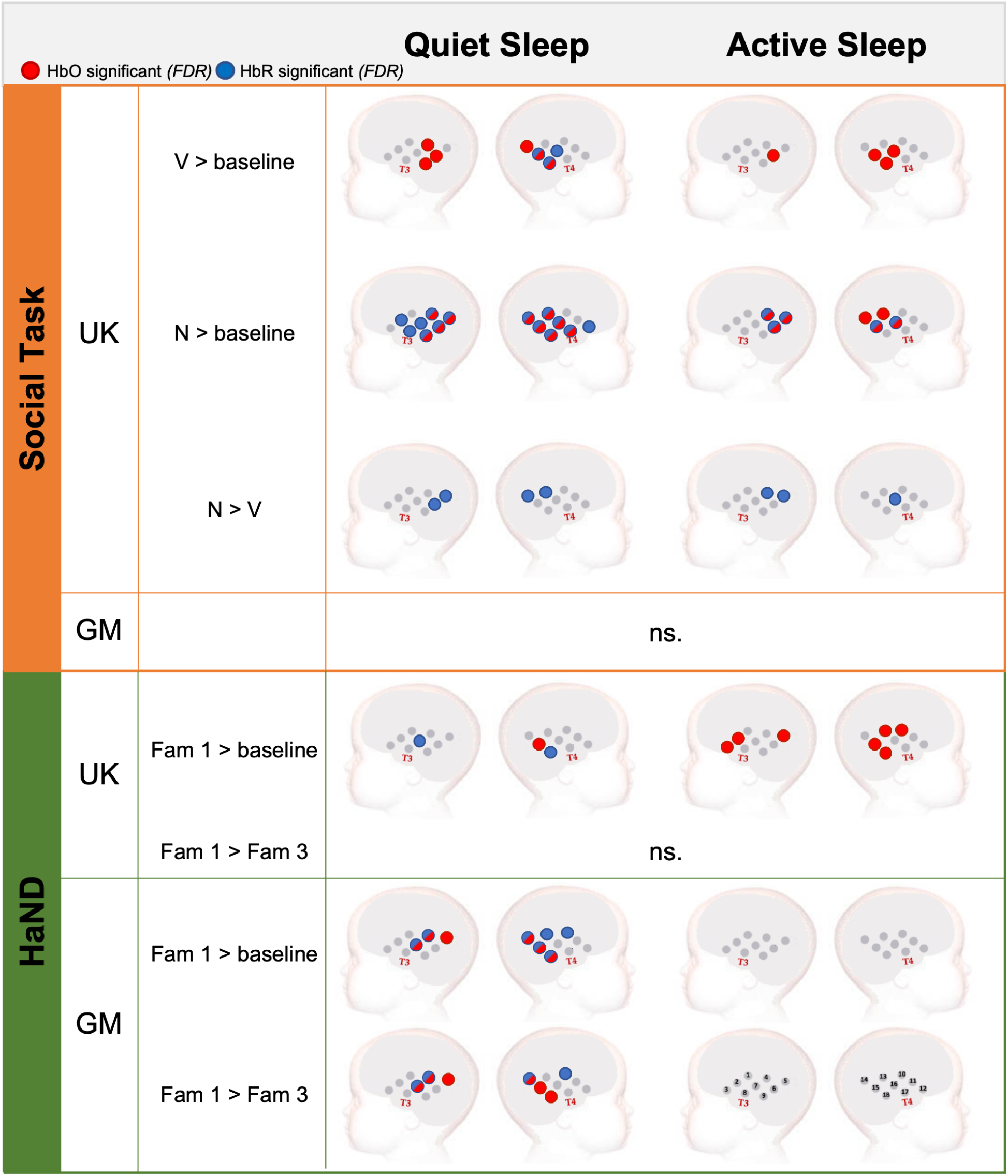
Results of the channel-wise analysis of hemodynamic activation for HbO (red) and HbR (blue) to stimuli of the social selectivity and habituation and novelty detection (HaND) paradigms, as well as condition contrasts, for the UK and Gambian (GM) cohorts of infants; ns - not significant.

**Table 3.** Deoxyhemoglobin (HbR) channels with significant non-vocal over vocal preference in quiet and active sleep stages, UK Cohort.

| Sleep State | Channel | Non-vocal Mean HbR (SD) | Vocal Mean HbR (SD) | *t* value | *p* (FDR-corrected) |
| --- | --- | --- | --- | --- | --- |
| QS | 5 | -0.15 (0.15) | -0.009 (0.12) | -4.8 | < 0.01** |
|  | 9 | -0.18 (0.11) | -0.04 (0.15) | -3.3 | 0.02* |
|  | 13 | -0.20 (0.19) | -0.03 (0.16) | -4.1 | < 0.01** |
|  | 14 | -0.19 (0.15) | -0.06 (0.16) | -2.9 | 0.03* |
| AS | 4 | -0.20 (0.25) | -0.02 (0.19) | -3.5 | 0.01* |
|  | 5 | -0.22 (0.25) | -0.05 (0.2) | -2.7 | 0.03* |
|  | 16 | -0.22 (0.33) | -0.06 (0.2) | -2.5 | 0.04* |

#### 3.4.2 Sleep-stage specific spatial distribution of significant response to the HaND paradigm

First, similarly to the social selectivity paradigm, we conducted channel-by-channel analysis of hemodynamic activation to Fam1 in AS and QS (one-sample t-test, signal *>* 0, FDR-corrected for the entire set of channels, *p adj <* 0.05) (Fig. 6). In the UK cohort, infants in QS showed activation in 1 out of 18 HbO channels (∼6%) and 2 HbR channels (∼11%); infants in AS showed activation in 7 HbO channels (∼39%) and 0 HbR channels. In the GM cohort, infants in QS showed activation in 6 HbO channels (∼33%) and 7 HbR channels (∼25%); infants in AS had no significant activation in HbO or HbR channels. Next, channels with significant activation to Fam1 were tested for habituation (Fam1 *>* Fam3) or dishabituation (Fam3 *>* Fam1) separately for each sleep stage group (paired-sample t-test, two-tailed, FDR-corrected for the set of channels significant against baseline, *p adj <* 0.05). In the UK cohort, we found that neither response was present in AS or QS. For the GM cohort, significant habituation response was found in 6 HbO channels (channels 4, 5, 7, 14, 15, 18) and 4 HbR channels (channels 4, 7, 10, 14) in the QS group. No channels with significant habituation response were identified in the AS group.

The results of the channel-by-channel analysis of hemodynamic activation to Novel trials in AS and QS (one-sample t-test, signal *>* 0, FDR-corrected for the entire set of channels, *p adj <* 0.05) identified no HbO or HbR channels with significant activation in either cohort. Therefore, we did not proceed with the analysis of sleep stage effect on the presence of novelty response (Novel *>* Fam3).

## 4 Discussion

This study examined the potential influence of sleep stage on infant brain hemodynamic responses across two auditory paradigms in UK and Gambian cohorts, using fNIRS. By leveraging both region-based and channel-wise analyses, we found that the effect of sleep stage on cortical processing was nuanced, depending on task type and population. The social selectivity paradigm results showed no sleep stage differences in overall amplitude or selectivity across conditions in either cohort. In the UK cohort, AS was associated with a higher HbO response during the first phase of the habituation paradigm (Fam1) and a significantly stronger habituation compared to QS. In contrast, in the Gambian cohort, QS was linked to more widespread activation and evidence of habituation compared to AS. Across both tasks, channel-wise analyses highlighted differences in the spatial distribution of activation by sleep stage. These results suggest that sleep stage may modulate early auditory processing in task- and population-specific ways, underscoring the need to understand its effect in infant neuroimaging research.

*Social selectivity paradigm.* The analyses based on the regions of interest defined in the full cohort of infants allowed us to examine the sleep stage effects on the amplitude of significant responses and the strength of response selectivity without being constrained by the low number of overlapping channels between sleep stage groups. We found that the amplitude of hemodynamic responses to the non-vocal and vocal conditions did not vary across sleep stages in either cohort of infants. Examining the hemodynamic response averaged over the ROI identified as having significant preference for non-vocal compared to vocal stimuli in the full cohort of infants by Greenhalgh et al.,^44^ we found the strength of non-vocal selectivity to be unaffected by sleep stage either for HbO or HbR signals. These finding aligns with a previous fNIRS study that found no sleep stage effect on response amplitude to speech and music stimuli in neonates^26^ and previous EEG studies that found no differences in the detection of change in auditory stimuli according to sleep stage in neonates.^12–14^

Channel-wise analyses using the sub-sample of infants with available sleep stage coding allowed us to examine the possible effect of sleep stage on the location of the response to social selectivity and habituation paradigms. In the UK cohort, infants in active and quiet sleep showed significant auditory response to both vocal and non-vocal stimuli in the medial and posterior temporal channels, with no apparent lateralization. The response to both types of stimuli was more widespread in quiet compared to active sleep with approximately double the number of active channels. Infants in both sleep stage groups discriminated between vocal and non-vocal stimuli with a significant selectivity for non-vocal stimuli, albeit only in the HbR signal. Despite wider activation in QS to both conditions, the number of channels with significant non-vocal preference was similar across sleep stage groups. The finding of non-vocal preference is in line with the results obtained using the full cohort of infants^44^ and with previous studies on this age group.^47, 58, 59^

In our sub-sample of the Gambian cohort, no channels showed significant activation to either condition relative to baseline, therefore, condition contrasts were not investigated. This partly contradicts the results obtained using the full cohort of infants where significant activation to both conditions was found, however, stimuli selectivity was also not present in the full cohort. Hence, while the small sample used in this analysis could account for the inconsistent results, the investigation of condition contrasts in this paradigm at group level would still not be possible even with a bigger sample. Interestingly, the full cohort paper also found a high level of inter-individual variability in social selectivity with some infants showing N *>* V and some V *>* N selectivity across both cohorts. Future work should explore this further as N - V discrimination also associated with FC maturation patterns.^44^

In summary, our ROI-based and channel-wise analyses suggest that infants in both active and quiet sleep stages respond robustly to auditory stimuli and are equally able to discriminate between vocal and non-vocal sounds. Sleep stage did not affect the amplitude, spatial distribution, or direction of stimulus selectivity. Importantly, while it is well established that non-vocal selectivity in early infancy typically shifts toward vocal preference with development, our results show that at one month of age, infants consistently exhibit non-vocal preference regardless of sleep stage.

This finding is a valuable contribution to the field because it suggests that core auditory discrimination processes and early perceptual biases are preserved across sleep stages in two different populations of infants. Furthermore, by showing that sleep stage does not confound early markers of auditory selectivity, this study helps refine methodological approaches in infant neuroimaging and strengthens developmental models of early auditory social processing.

*Habituation and novelty detection.* Results based on the ROI defined by Greenhalgh et al.^44^ showed a higher average amplitude of the hemodynamic response to the Fam1 epoch in AS compared to QS in the UK cohort. Infants in AS also showed significantly stronger habituation, with an average HbO concentration decrease of 0.49*µ*Mol between first and last familiarization epochs. In contrast, infants that remained in QS showed a mean difference of -0.12*µ*Mol between Fam1 and Fam3, indicating an absence of habituation in QS.

Channel-wise analysis further supported greater response to the HaND stimuli in AS in the UK cohort, with more channels showing significant activation to Fam1 compared to QS. In AS, significant responses were found in bilateral frontal and posterior temporal regions, while in QS, activation was limited to posterior temporal regions. However, in the AS group, the activation was observed only in the HbO signal, whereas in QS, both HbO and HbR signals showed significant responses. In contrast to ROI-based analysis, no significant habituation patterns (channel-level differences between Fam1 and Fam3 epochs) were found in either sleep stage group. This contrasts with findings from the full UK cohort, where habituation effects were evident, suggesting that reduced power in our subsample may underlie the lack of significant habituation.

These findings are in line with Kotilahti et al. (2005), who reported a stronger hemodynamic response to sinusoidal tones in AS compared to QS.^25^ However, their later study using more complex auditory stimuli (speech and music) found no effect of sleep stage on the strength of the response to either stimuli.^26^ Importantly, however, both studies involved newborns considerably younger than the participants in the present study. In an investigation with slightly older infants, Hunter and colleagues^19^ also found a decrease in response between subsequent presentations of auditory stimuli in AS but not QS, consistent with our UK results.

In the Gambian cohort, ROI-based analysis revealed no differences in average response amplitude to Fam1 across sleep stages. The strength of habituation was also not affected by sleep stage. However, according to the results of the channel-wise analysis, channels with significant activation to Fam1, as well as significant habituation response (Fam1 *>* Fam3) were found only in the QS group, while the response in AS did not differ from baseline in any of the channels. This contrasts with the results of the UK cohort and prior findings of stronger habituation in AS.^18, 19^

The discrepancy may stem from the smaller sample size, higher inter-individual variability, and slightly lower data quality in the AS group in the Gambian sample. Alternatively, activation in QS but not AS may reflect a population-specific hemodynamic response profile.

In summary, this analysis reveals that sleep stage modulates hemodynamic responses to repeated auditory stimuli in one-month-old infants, with different patterns emerging across UK and Gambian cohorts. In the UK group, AS was associated with stronger initial responses and evidence of habituation, whereas in the Gambian group, QS showed more robust activation. These findings highlight variability in early sensory processing and emphasize the importance of accounting for sleep stage when interpreting infant neuroimaging data.

The presence of a sleep stage effect on the amplitude of the response to the habituation paradigm, but not to the social selectivity paradigm, may be partly explained by differences in stimulus characteristics. The habituation paradigm used a relatively long, continuous 8-second spoken sentence, which likely engages sustained auditory and linguistic processing and may thus be more sensitive to cortical state fluctuations across sleep stages. In contrast, the social selectivity paradigm consisted of short, non-speech vocalizations (such as laughing, crying, yawning) and various environmental sounds (toys, rattles, bells), which are acoustically diverse and likely elicit hemodynamic responses that are more directly driven by the sensory properties of the stimuli - that is, processed in a more stimulus-dependent, bottom-up manner, less reliant on the modulatory influence of sleep stage. Furthermore, spoken sentences have exaggerated prosody and pitch contours designed to capture infant attention, possibly amplifying differences between active and quiet sleep due to differences in cortical excitability or sensory gating across sleep stages. Together, these differences in stimulus duration, complexity, and linguistic content may explain why sleep stage modulated responses in the habituation paradigm but not in the social selectivity paradigm.

The differences between our results and previous studies could stem from several methodological and developmental factors. First, our study included slightly older infants than some of the earlier work, which may have influenced how sleep stages modulate hemodynamic responses. As active and quiet sleep mature into REM and NREM sleep, respectively, these stages become more differentiated and structured, with distinct physiological and neural profiles. For example, sleep spindles, are absent at birth but develop rapidly within the first months of life, marking a transition from quiet sleep to NREM sleep. This changing sleep architecture means that the functional significance of sleep stages and their influence on sensory processing may vary substantially with age. Second, differences in stimulus type could also contribute to differences in findings, although our results indicate this relationship is not straightforward. In our study, the more complex, speech-like stimuli of the habituation paradigm showed sleep stage differences in response amplitude, while the shorter non-speech sounds of the social selectivity paradigm did not, yet quiet sleep still recruited more channels overall in the latter. This indicates that the influence of sleep stage may not simply track stimulus complexity, but rather reflect interactions between stimulus features, developmental stage, and the neural networks engaged. Third, variations in sample size and resulting statistical power likely contribute, especially as our subsample stratified by sleep stage was considerably smaller than our full cohort analyses, potentially reducing sensitivity to detect habituation or condition contrasts. Finally, inconsistencies may also arise due to differences in the neuroimaging modalities themselves; while fNIRS measures slow hemodynamic changes, MEG and EEG capture rapid electrophysiological dynamics, which may be differentially sensitive to sleep stage. Together, these factors highlight the multifaceted nature of sleep stage effects on early auditory processing and underscore the need for larger studies with different populations of infants to fully disentangle how sleep modulates infant brain responses.

### 4.1 Strengths and limitations

One limitation of this study is the observed variation in data quality between sleep stages, which may have influenced the precision of our findings. Specifically, data quality and the number of valid trials were generally lower during active sleep compared to quiet sleep across both cohorts, with the exception of the number of vocal trials in the UK cohort, which did not differ significantly by sleep stage. Nevertheless, it is important to highlight that despite these differences, the proportion of clean data remained consistently high (above 95% across all conditions), indicating that while data acquisition may pose more challenges during active sleep, the quality was still sufficient to ensure the reliability of our analyses. Another limitation relates to sample size. Although adequate for detecting general patterns, our stratified subsample did not allow for more sensitive analyses such as TFCE, which were feasible in the full cohort. As a result, we were unable to examine whether the time window of the peak hemodynamic response differed by sleep stage - a factor that could be relevant, since differences in response latency might underlie some of the observed variability between stages. Finally, our reliance on behavioral sleep stage classification, based on the Brazelton Neonatal Behavioral Assessment Scale^5^ and the Anders Manual,^54^ represents a limitation. Although this method is practical and widely used in infant research, it lacks the precision of gold-standard multimodal approaches that combine EEG, ECG, respiration, and behavioral scoring for more accurate sleep stage coding.

Additionally, our sample size was reduced due to the poor quality of some video recordings or key behaviors and respiration cues being obscured - often due to how infants were held or wrapped during testing. This issue is not unique to our study; it is a known challenge in naturalistic infant research, where standardizing infant positioning and camera angles is difficult. In The Gambia, it is customary to wrap infants in multiple layers of fabric - making visual assessment of breathing and subtle movements particularly difficult.

These challenges highlight the value of developing automated methods for sleep stage classification based solely on neural, hemodynamic, or cardiac signals. Such approaches would not only improve the accuracy and objectivity of sleep staging but would also significantly reduce the need for time-intensive behavioral coding and specialized training, thereby enhancing the scalability of sleep-related research in infancy.

### 4.2 Conclusions

Our study offers several unique contributions to the field of developmental cognitive neuroscience and sleep research. First, our study stands out by investigating sleep stage effects in two populations of infants, which strengthens our findings by allowing us to examine the consistency effects across different samples. Furthermore, we investigated sleep stage effects across two distinct auditory paradigms - social selectivity and habituation and novelty detection. Lastly, we employed two complementary analytical approaches: one guided by regions of interest defined from hemodynamic responses in the full cohort to maximize signal-to-noise ratio, and another channel-wise approach that allowed us to explore potential sleep stage effects on the spatial distribution and presence of brain activation.

In conclusion, this study provides new insights into how sleep stages may shape early brain responses to auditory processing across two distinct populations and paradigms. The results of the study highlight the nuanced role of sleep stage in shaping hemodynamic responses during early infancy. These findings underscore the importance of accounting for sleep stages in infant neuroimaging research and point to the need for larger studies with different populations to further unravel how sleep architecture interacts with early sensory and cognitive development.

## Acknowledgments

We would like to thank the families who took part in this study. Furthermore we would like to thank the wider BRIGHT Project team both in the Gambia and the UK, without whom this work would not be possible. The BRIGHT Study is funded by the Gates Foundation (OPP1127625 and core funding MC-A760-5QX00 to the International Nutrition Group by the Medical Research Council UK and the UK Department for International Development (DfID) under the MRC/DfID COn-cordant agreement. S.L.F is supported by a UKRI Future Leaders Fellowship (MR/S018425/1).

S.E.M is supported by a Wellcome Trust Senior Research Fellowship (220225/Z/20/Z).

*The funders had no role in study design, data collection and analysis, decision to publish, or preparation of the manuscript*.

## References

1 L. I. van Dyck and E. M. Morrow, “Genetic control of postnatal human brain growth,” Current opinion in neurology 30(1), 114–124 (2017). 10.1097/WCO.0000000000000405.

2 J. Gervain, Y. Minagawa, L. Emberson, et al., “Using functional near-infrared spectroscopy to study the early developing brain: future directions and new challenges,” Neurophotonics 10(2), 023519–023519 (2023).

3 W.-L. Chen, J. Wagner, N. Heugel, et al., “Functional near-infrared spectroscopy and its clinical application in the field of neuroscience: advances and future directions,” Frontiers in neuroscience 14, 724 (2020).

4 M. S. Knoop, E. R. de Groot, and J. Dudink, “Current ideas about the roles of rapid eye movement and non–rapid eye movement sleep in brain development,” Acta Paediatrica 110(1), 36–44 (2021).

5. 5 T. B. Brazelton, Neonatal behavioral assessment scale / T. Berry Brazelton and J. Kevin Nugent., Clinics in developmental medicine ; no. 190, Mac Keith Press, London, 4th ed. ed. (2011).

6 M. Grigg-Damberger, D. Gozal, C. L. Marcus, et al., “The visual scoring of sleep and arousal in infants and children,” Journal of Clinical Sleep Medicine 3(02), 201–240 (2007).

7 M. Chipaux, M. T. Colonnese, A. Mauguen, et al., “Auditory stimuli mimicking ambient sounds drive temporal “delta-brushes” in premature infants,” PloS one 8(11), e79028 (2013).

8 S. Lori, S. Gabbanini, M. Bastianelli, et al., “Multimodal neurophysiological monitoring in healthy infants born at term: normative continuous somatosensory evoked potentials data,” Developmental Medicine & Child Neurology 59(9), 959–964 (2017).

9 P. Nevalainen, E. Pihko, M. Metsäranta, et al., “Evoked magnetic fields from primary and secondary somatosensory cortices: a reliable tool for assessment of cortical processing in the neonatal period,” Clinical neurophysiology 123(12), 2377–2383 (2012).

10 E. Pihko, A. Sambeth, P. Leppänen, et al., “Auditory evoked magnetic fields to speech stimuli in newborns–effect of sleep stages,” Neurol. Clin. Neurophysiol 6, 1–5 (2004).

11 A. Sambeth, K. Ruohio, P. Alku, et al., “Sleeping newborns extract prosody from continuous speech,” Clinical Neurophysiology 119(2), 332–341 (2008).

12 A. Sambeth, S. Pakarinen, K. Ruohio, et al., “Change detection in newborns using a multiple deviant paradigm: a study using magnetoencephalography,” Clinical Neurophysiology 120(3), 530–538 (2009).

13 O. Martynova, J. Kirjavainen, and M. Cheour, “Mismatch negativity and late discriminative negativity in sleeping human newborns,” Neuroscience letters 340(2), 75–78 (2003).

14 K. M. Uhler, S. K. Hunter, E. Tierney, et al., “The relationship between mismatch response and the acoustic change complex in normal hearing infants,” Clinical Neurophysiology 129(6), 1148–1160 (2018).

15 M. Cheour, R. Čėponiené, P. Leppänen, et al., “The auditory sensory memory trace decays rapidlyin newborns,” Scandinavian Journal of Psychology 43(1), 33–39 (2002).

16 K. Wanrooij, P. Boersma, and T. L. Van Zuijen, “Fast phonetic learning occurs already in 2-to-3-month old infants: an erp study,” Frontiers in psychology 5, 77 (2014).

17 T. van Leeuwen, P. Been, M. van Herten, et al., “Two-month-old infants at risk for dyslexia do not discriminate/bak/from/dak: A brain-mapping study,” Journal of neurolinguistics 21(4), 333–348 (2008).

18 F. McNamara, H. Wulbrand, and B. T. Thach, “Habituation of the infant arousal response,” Sleep 22(3), 320–326 (1999).

19 S. K. Hunter, S. J. Gillow, and R. G. Ross, “Stability of p50 auditory sensory gating during sleep from infancy to 4 years of age,” Brain and Cognition 94, 4–9 (2015).

20 B. Blanco, M. Molnar, I. Arrieta, et al., “Functional brain adaptations during speech processing in 4-month-old bilingual infants,” Developmental Science 28(1), e13572 (2025).

21 P. Zaramella, F. Freato, A. Amigoni, et al., “Brain auditory activation measured by near-infrared spectroscopy (nirs) in neonates,” Pediatric research 49(2), 213–219 (2001).

22 M. Pena, A. Maki, D. Kovačić, et al., “Sounds and silence: an optical topography study of language recognition at birth,” Proceedings of the National Academy of Sciences 100(20), 11702–11705 (2003).

23 Y. Saito, S. Aoyama, T. Kondo, et al., “Frontal cerebral blood flow change associated with infant-directed speech,” Archives of Disease in Childhood-Fetal and Neonatal Edition 92(2), F113–F116 (2007).

24 Y. Saito, T. Kondo, S. Aoyama, et al., “The function of the frontal lobe in neonates for response to a prosodic voice,” Early human development 83(4), 225–230 (2007).

25 K. Kotilahti, I. Nissilä, M. Huotilainen, et al., “Bilateral hemodynamic responses to auditory stimulation in newborn infants,” Neuroreport 16(12), 1373–1377 (2005).

26 K. Kotilahti, I. Nissilä, T. Näsi, et al., “Hemodynamic responses to speech and music in newborn infants,” Human brain mapping 31(4), 595–603 (2010).

27 T. Arimitsu, Y. Minagawa, T. Yagihashi, et al., “The cerebral hemodynamic response to phonetic changes of speech in preterm and term infants: The impact of postmenstrual age,” NeuroImage: Clinical 19, 599–606 (2018).

28 O. W. Lee, D. Mao, J. Wunderlich, et al., “Two independent response mechanisms to auditory stimuli measured with functional near-infrared spectroscopy in sleeping infants,” Trends in Hearing 28, 23312165241258056 (2024).

29 F. Homae, H. Watanabe, T. Nakano, et al., “The right hemisphere of sleeping infant perceives sentential prosody,” Neuroscience research 54(4), 276–280 (2006).

30 T. Nakano, H. Watanabe, F. Homae, et al., “Prefrontal cortical involvement in young infants’ analysis of novelty,” Cerebral Cortex 19(2), 455–463 (2009).

31 G. Taga, H. Watanabe, and F. Homae, “Developmental changes in cortical sensory processing during wakefulness and sleep,” NeuroImage 178, 519–530 (2018).

32 D. Mao, J. Wunderlich, B. Savkovic, et al., “Speech token detection and discrimination in individual infants using functional near-infrared spectroscopy,” Scientific reports 11(1), 24006 (2021).

33 O. W. Lee, D. Gao, T. Peng, et al., “Measuring speech discrimination ability in sleeping infants using fnirs—a proof of principle,” Trends in Hearing 29, 23312165241311721 (2025).

34 H. Shuto, A. Yasuhara, T. Sugimoto, et al., “Longitudinal determination of cerebral blood flow velocity in neonates with the doppler technique,” Neuropediatrics 18(04), 218–221 (1987).

35 G. Jorch, T. Huster, and H. Rabe, “Dependency of doppler parameters in the anterior cerebral artery on behavioural states in preterm and term neonates,” Neonatology 58(2), 79–86 (1990).

36 T. Näsi, J. Virtanen, T. Noponen, et al., “Spontaneous hemodynamic oscillations during human sleep and sleep stage transitions characterized with near-infrared spectroscopy,” PLoS One 6(10), e25415 (2011).

37 D. M. Münger, H. U. Bucher, and G. Duc, “Sleep state changes associated with cerebral blood volume changes in healthy term newborn infants,” Early human development 52(1), 27–42 (1998).

38 F. Y. Wong, N. B. Witcombe, S. R. Yiallourou, et al., “Cerebral oxygenation is depressed during sleep in healthy term infants when they sleep prone,” Pediatrics 127(3), e558–e565 (2011).

39 A. Tokariev, J. A. Roberts, A. Zalesky, et al., “Large-scale brain modes reorganize between infant sleep states and carry prognostic information for preterms,” Nature communications 10(1), 2619 (2019).

40 C. W. Lee, B. Blanco, L. Dempsey, et al., “Sleep state modulates resting-state functional connectivity in neonates,” Frontiers in Neuroscience 14, 347 (2020).

41 J. Uchitel, B. Blanco, L. Collins-Jones, et al., “Cot-side imaging of functional connectivity in the developing brain during sleep using wearable high-density diffuse optical tomography,” NeuroImage 265, 119784 (2023).

42 I. Mueller, R. X. Rodriguez, N. Pini, et al., “Infant sleep state coded from respiration and its relationship to the developing functional connectome: a feasibility study,” Developmental Cognitive Neuroscience 72, 101525 (2025).

43 S. Lloyd-Fox, S. McCann, B. Milosavljevic, et al., “The brain imaging for global health (bright) project: longitudinal cohort study protocol,” Gates Open Research 7, 126 (2024).

44. 44 I. Greenhalgh, B. Blanco, C. Bulgarelli, et al., “Cross-paradigm fnirs brain activity in neonates across the gambia and uk,” Biorxiv (2025).

45 S. Lloyd-Fox, A. Blasi, S. McCann, et al., “Habituation and novelty detection fnirs brain responses in 5-and 8-month-old infants: The gambia and uk,” Developmental Science 22(5), e12817 (2019).

46 E. J. Jones, L. Mason, J. B. Ali, et al., “Eurosibs: Towards robust measurement of infant neurocognitive predictors of autism across europe,” Infant Behavior and Development 57, 101316 (2019).

47 S. Lloyd-Fox, A. Blasi, E. Mercure, et al., “The emergence of cerebral specialization for the human voice over the first months of life,” Social Neuroscience 7(3), 317–330 (2012).

48 S. Lloyd-Fox, A. Blasi, C. Elwell, et al., “Reduced neural sensitivity to social stimuli in infants at risk for autism,” Proceedings of the Royal Society B: Biological Sciences 280(1758), 20123026 (2013). 10.1098/rspb.2012.3026.

49 S. Lloyd-Fox, C. W. Lee, L. Pirazzoli, et al., “Establishing a biomarker of cortical specialisation in infants in the gambia and uk,” The FASEB Journal 30, 1149–14 (2016). 10.1096/fasebj.30.1_supplement.1149.14.

50 N. Everdell, A. Gibson, I. Tullis, et al., “A frequency multiplexed near-infrared topography system for imaging functional activation in the brain,” Review of Scientific Instruments 76(9) (2005).

51 S. Lloyd-Fox, J. E. Richards, A. Blasi, et al., “Coregistering functional near-infrared spectroscopy with underlying cortical areas in infants,” Neurophotonics 1(2), 025006–025006 (2014).

52. 52 S. M. Hernandez and L. Pollonini, “Nirsplot: a tool for quality assessment of fnirs scans,” in Optics and the Brain, BM2C–5, Optica Publishing Group (2020).

53 T. J. Huppert, S. G. Diamond, M. A. Franceschini, et al., “Homer: a review of time-series analysis methods for near-infrared spectroscopy of the brain,” Applied optics 48(10), D280– D298 (2009).

54. 54 T. Anders, R. Emde, and A. Parmelee Jr, “A manual ofstandardized terminology, techniques and criteria for the scoring ofstates ofsleep and wakefulness in newborn infants,” (1971).

55 R Core Team, R: A Language and Environment for Statistical Computing. R Foundation for Statistical Computing, Vienna, Austria (2024).

56 Y. Benjamini and Y. Hochberg, “Controlling the false discovery rate: a practical and powerful approach to multiple testing,” Journal of the Royal statistical society: series B (Methodological) 57(1), 289–300 (1995).

57 S. M. Smith and T. E. Nichols, “Threshold-free cluster enhancement: addressing problems of smoothing, threshold dependence and localisation in cluster inference,” Neuroimage 44(1), 83–98 (2009).

58 T. Grossmann, R. Oberecker, S. P. Koch, et al., “The developmental origins of voice processing in the human brain,” Neuron 65(6), 852–858 (2010).

59 A. Blasi, E. Mercure, S. Lloyd-Fox, et al., “Early specialization for voice and emotion processing in the infant brain,” Current biology 21(14), 1220–1224 (2011).

