## Supplemental Materials for "Effect of sleep stages on patterns of fNIRS hemodynamic response to auditory paradigms in one-month-old Gambian and UK infants"

### 1 Supplemental Materials

#### 1.1 Confidence levels of sleep stage classification criteria

Supplemental Table 1 provides an overview of the criteria used for sleep stage classification and their associated confidence levels. Each criterion's percentage reflects its relative contribution to determining the sleep stage. For instance, 'Eyes closed/ Eyes Open' and 'Rapid Eye Movements (REM) versus Non-REM (NREM)' have the highest confidence levels at 25%, indicating their significant role in classification between the two stages. If any of the criteria can not be coded due to obscured visibility, the confidence level decreases accordingly.

**Table 1** Overview of The Sleep Classification Criteria and their Corresponding Level of Confidence.

| **Criterium** | **Confidence (%)** |
|---|---|
| *Eyes closed versus Eyes Open* | 25 |
| *REM versus NREM* | 25 |
| *Regular versus Irregular breathing* | 15 |
| *Jerky versus Smooth movement* | 15 |
| *Sucking* | 5 |
| *Facial Grimace* | 5 |
| *Vocalization* | 5 |
| *Startles* | 5 |

#### 1.2 Retention of participants based on fNIRS quality control

Supplemental Figure 1 shows participant retention per paradigm based on fNIRS data quality in the two cohorts. The data used in the current analysis was pre-processed as part of a separate investigation using the full dataset (Greenhalgh et al., 2025). In the Gambian (GM) cohort, of the total number of participants eligible at one month ($N$ = 204), 136 had valid fNIRS data for at least the habituation part (Fam1, Fam2, Fam3) of the Habituation and Novelty Detection (HaND) paradigm and 148 had valid fNIRS data for the social selectivity paradigm. In the UK cohort, of the total number of participants eligible at one month ($N$ = 61), 46 had valid fNIRS data for the social selectivity paradigm and 38 for the HaND paradigm. Data collected as part of the functional connectivity (FC) paradigm was analysed as part of the separate full cohort investigation, but due to the small number of participants with valid FC and sleep stage coding, the effect of sleep stages on FC patterns was not investigated as part of the current work.

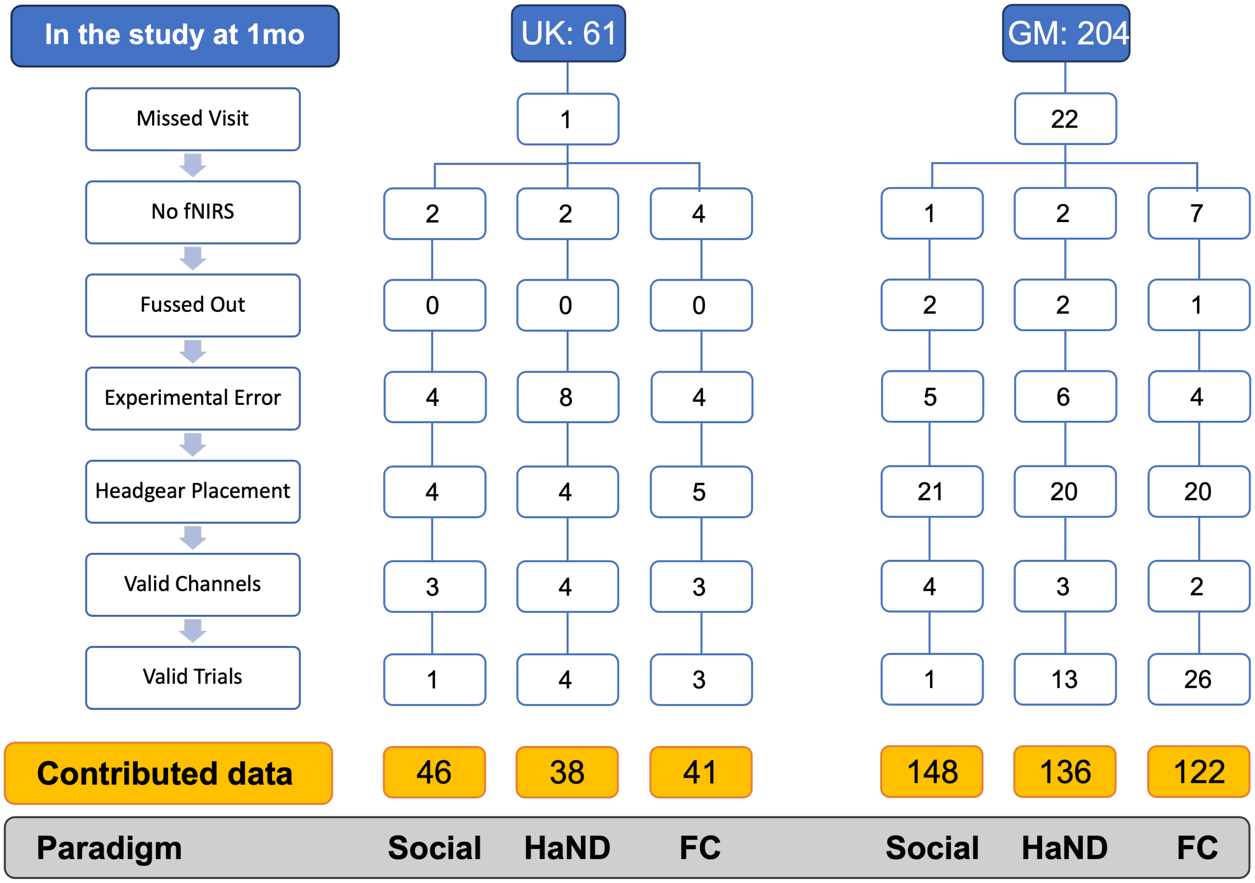


**Fig 1** Reasons for exclusion of participants from the fNIRS analysis from the United Kingdom (UK) and Gambian (GM) cohorts across the three paradigms: vocal/non-vocal selectivity task (Social), Habituation and Novelty Detection (HaND) and functional connectivity (FC); The reasons for exclusion were: Missed Visit - participant did not attend the one month session; No fNIRS - participant attended the one month session, but fNIRS data was not collected at all or for specific task; Fussed Out - infant woke up and cried during scanning so the recording had to be stopped; Experimental Error - collected data is invalid due to missing photos of the headband placement, missing event markers or technical issues; Headgear Placement - collected data is invalid due to wrong placement of the fNIRS headband; Valid Channels - collected data is invalid due to too few valid channels; Valid Trials - collected data is invalid due to too few valid trials.

##### *1.3 Effect of sleep stages on the amplitude of significant HbR responses*

Hemodynamic responses at one month of age are typically reduced in magnitude compared to those observed in older children and adults, presenting challenges for interpreting hemodynamic activation. Among the two chromophores, oxyhemoglobin (HbO) generally yields stronger task-related signals due to its higher signal-to-noise ratio, making it the preferred measure in infant fNIRS research (Lloyd-Fox et al., 2010; Gervain et al., 2011). Consequently, the main analyses in the primary manuscript focused on HbO. Nonetheless, given evidence suggesting that deoxyhemoglobin (HbR) may offer greater spatial specificity under certain conditions, we also examined HbR responses. These Supplemental analyses employed identical preprocessing and statistical procedures to those used for HbO and are reported here for completeness.

Supplemental Figure 2 presents the results of ROI-based analyses of the comparison of the mean amplitude of significant HbR responses to vocal and non-vocal conditions of the social selectivity paradigm between sleep stage groups. We found no significant differences in the mean response amplitude by sleep stage for any of the conditions in the UK or GM cohorts (all $p > 0.46$).

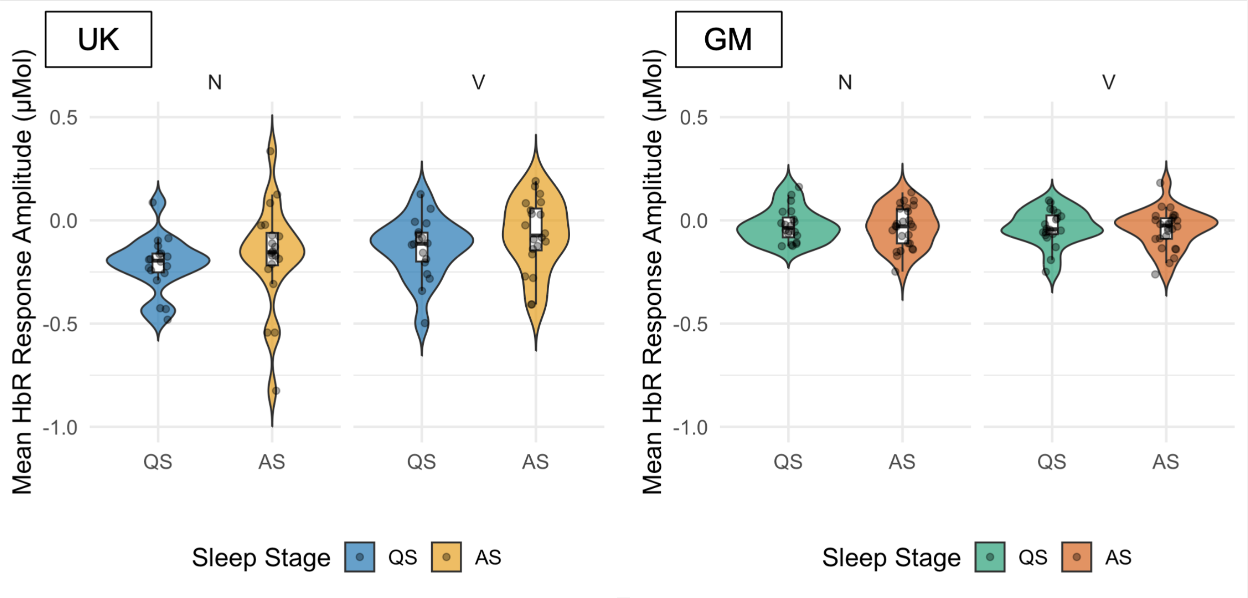


**Fig 2** The distribution of mean deoxyhemoglobin (HbR) response amplitudes ($\mu$Mol) to non-vocal (N) and vocal (V) trials of the social selectivity task across sleep stages (QS = Quiet Sleep, AS = Active Sleep) in the UK (left) and Gambian (GM) (right) cohorts. Each plot illustrates individual data points, overall distributions, and embedded boxplots.

Supplemental Figure 3 presents the results of ROI-based analyses of the comparison of the mean amplitude of significant HbR responses to first five familiarization trials (Fam1) of the HaND paradigm between sleep stage groups. While infants in AS showed a higher amplitude HbO response compared to QS infants in the UK cohort, HbR response was not significantly different between the sleep stage groups in either cohort (all $p > 0.51$).

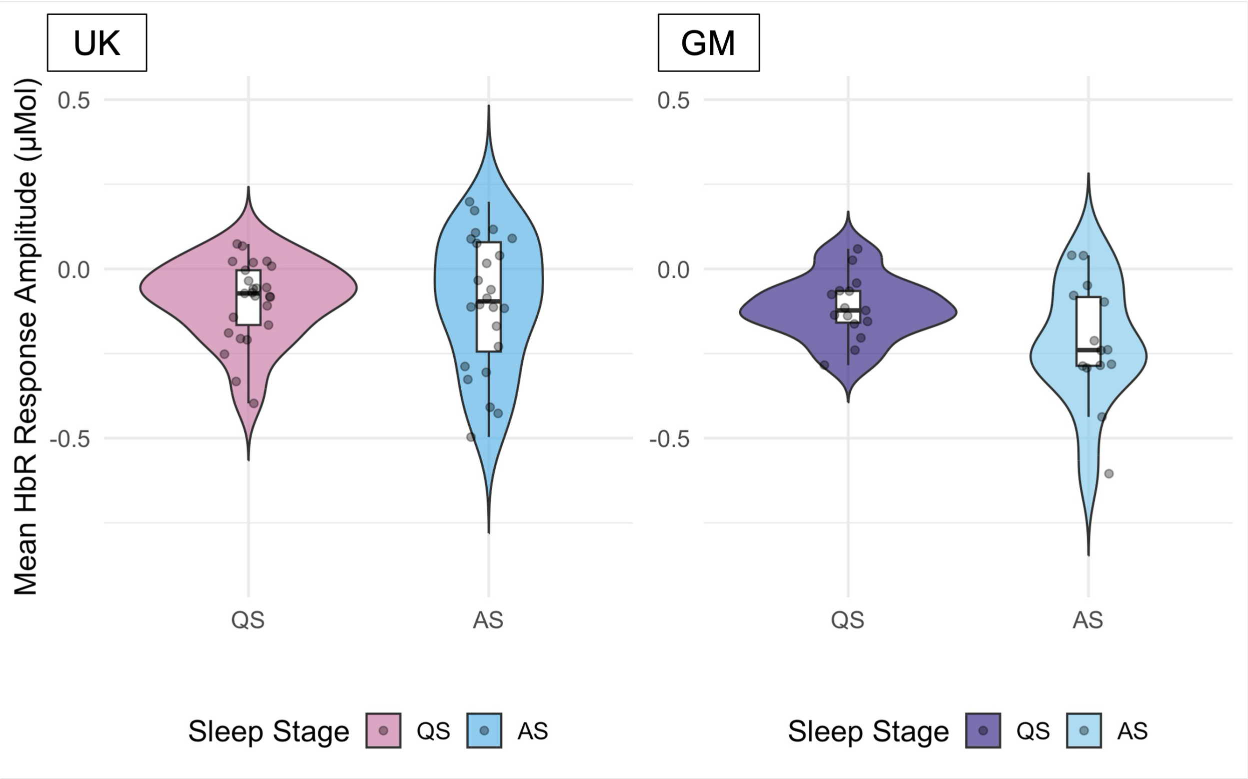


**Fig 3** The distribution of mean deoxyhemoglobin (HbR) response amplitudes ($\mu$Mol) to first five familiarization trials (Fam1) of the habituation and novelty detection (HaND) task across sleep states (QS = Quiet Sleep, AS = Active Sleep) in the UK (right) and Gambian (GM) (left) cohorts.

##### *1.4 The effect of sleep stages on the strength of non-vocal selectivity*

Supplemental Figure 4 presents the results of the ROI-based analyses that assessed whether sleep stage influenced the strength of non-vocal selectivity - that is, the degree to which infants responded more strongly to non-vocal than vocal stimuli. In the UK cohort, significant N $>$ V selectivity was identified in the full cohort analysis (Greenhalg et al., 2025). In the GM cohort, no N $>$ V or V $>$ N selective regions were observed. Therefore, for the current work, ROI-based analyses of the effect of sleep stage on the strength of N $>$ V selectivity were focused on the UK cohort. A two-sample t-test comparing average N–V condition contrast values between sleep stage groups revealed no significant difference for HbO or HbR in the UK cohort.

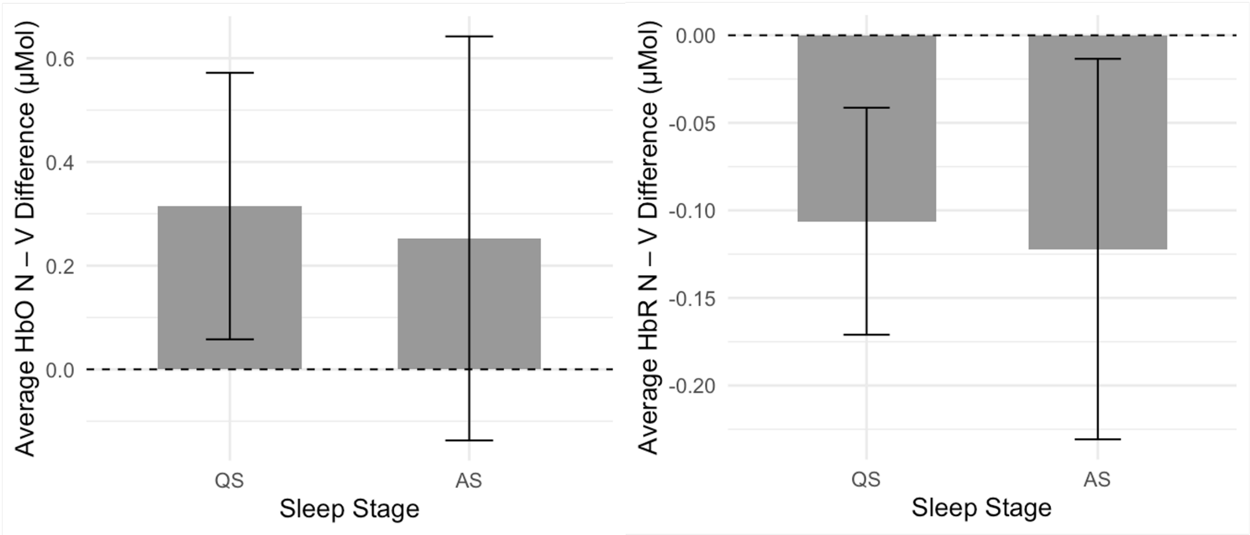


**Fig 4** Mean difference in oxyhemoglobin (HbO) (left) and deoxyhemoglobin (HbR) (right) concentration between Non-vocal (N) and Vocal (V) trials during the social selectivity task, shown separately for Quiet Sleep (QS) and Active Sleep (AS) for the UK cohort. Bars represent group means; error bars indicate 95% confidence intervals. For HbO, positive values reflect a greater HbO increase (i.e. stronger response) to N compared to V stimuli. For HbR, a negative value reflects a stronger response to Non-vocal compared to Vocal stimuli.

##### *1.5 The effect of sleep stages on the strength of HbR response habituation*

Supplemental Figure 5 presents the results of the ROI-based analyses that assessed whether sleep stage influenced the strength of habituation - that is, the degree to which infants responded more strongly to the first five familiarisation trials (Fam1) compared to the last five familiarisation trials (Fam3) of the HaND paradigm. While infants in AS showed greater habituation in their HbO response compared to QS infants in the UK cohort, HbR response was not significantly different between the sleep stage groups in either cohort (all $p > 0.16$).

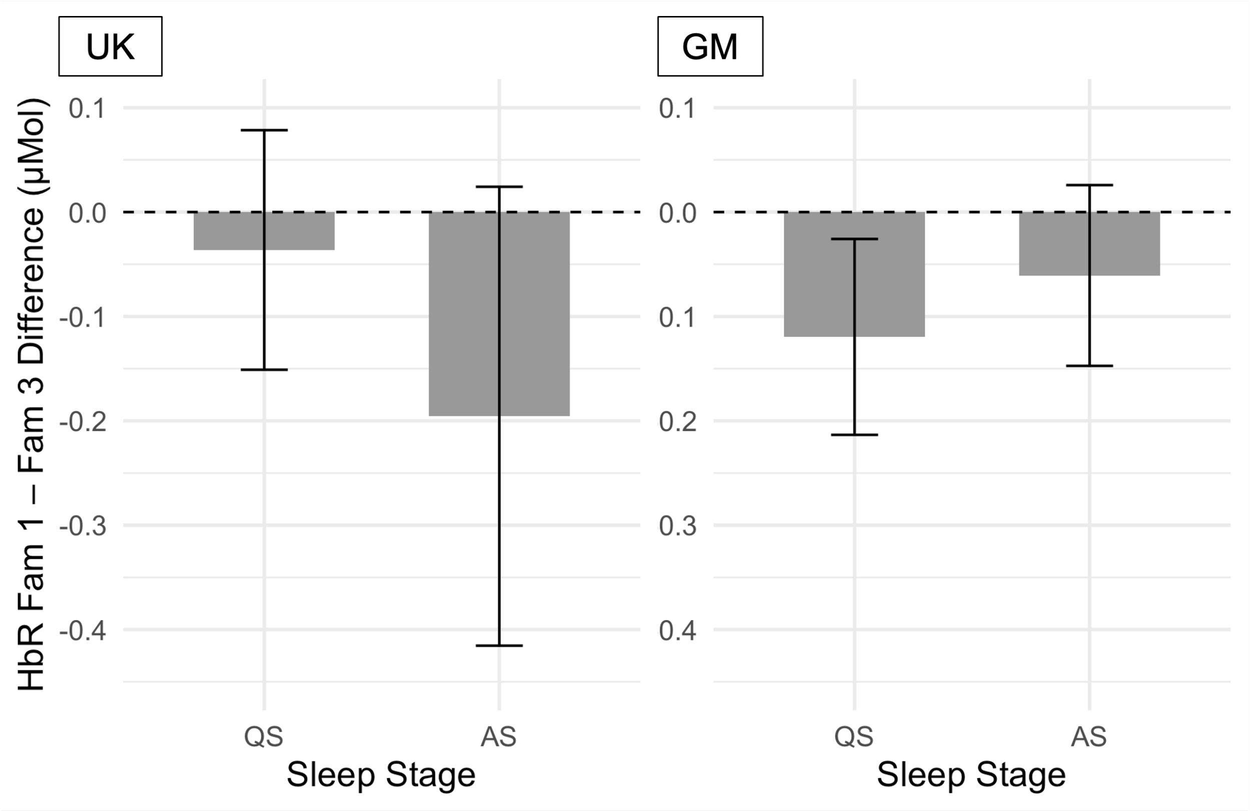


**Fig 5** Mean difference (Fam1 - Fam3) in deoxyhemoglobin (HbR) concentration by sleep stage in the UK (right) and Gambian (GM) (left) cohort. Error bars indicate standard errors.
